# In silico investigation of Y220C mutant p53 for lead design

**DOI:** 10.1101/820761

**Authors:** Mayank Roy Chowdhury, Anamika Tiwari, G.P. Dubey

**Affiliations:** Project assistant, Advance centre for traditional and Genomic Medicine, Institute of Medical Sciences, Banaras Hindu University, Varanasi-221 005; Research Scholar, Department of Kriya Sharir, Institute of Medical Sciences, Banaras Hindu University, Varanasi-221 005; Distinguished Professor, Department of Kriya Sharir, Institute of Medical Sciences, Banaras Hindu University, Varanasi-221 005

**Keywords:** Anticancer, docking, fragment-based design, molecular dynamics simulations, Y220C

## Abstract

p53 protein coded by the Tp53 gene is considered as one of the most intensively researched protein and mainly due to its role as a tumor suppressor, it acts as a tumor suppressor by carrying out two biologically complex processes namely Cell cycle arrest and apoptosis, In the oncogenic Y220C mutant p53, tyrosine is replaced by cysteine at 220^th^ residue of the DNA binding Domain which causes the formation of a surface crevice, this specific mutation is responsible for approx. 100,000 cancer cases per year due to the destabilization and denaturation of the protein, as a result, the protein degrades at room temperature. In this work we carry out intensive Molecular Dynamic Simulations and Molecular Docking Studies to understand the structural dynamics of wild type p53 and changes the occurs in the mutant protein and also try to design lead against the druggable crevice and at the end of our study we used fragment-based optimization to come up with lead molecules which can act as scaffold for further drug development process

## 1. Introduction

p53 protein is a homotetramer made of 393 amino acid residues, the name p53 refers to its apparent molecular mass, as it runs as a 53 kDa molecule on SDS-PAGE. The human p53 gene is located on the seventeenth chromosome, it’s a phosphoprotein made of 393 amino acids. The N-terminal region (residues 1-62) contains the transactivation domain, which is further divided into 2 subdomains and is followed by a proline-rich region (residues 63-64) important for apoptotic activity, The DBD also known as core domain (residues 94-292) is the domain for binding to the specific sequence on DNA. The nuclear localization signaling domain (residues 316-325) is involved in the intracellular localization of p53,The OD (residues 326-356) is responsible for tetramerization which is essential for the p53 activity, C-terminal regulatory domain CTD (residues 363-393) acts as a flexible region, and is involved in the downregulation of the central DBD (Wang et al. 1995).

The core domain is of 25-KD is inherently unstable and melts just above body temperature, based on NMR determined structure analysis it was found, the presence of many polar residues in the hydrophobic core which affects the dynamics and can be the reason behind its instability (Cañadillas et al. 2006). p53 core domain is a β-sandwich composed of 2 antiparallel β-sheets with a small β-hairpin (124-135) in contact with the second β-sheet, closing the access to the hydrophobic core,β-sandwich domains are subdivided in a variety of different folds, form immunoglobulin type fold found in antibodies. There are 2 more helices in the middle of loop2: one involved in Zn coordination (177-181) and a small 310 helix (166-168) Together, these structural elements form an extended surface that makes specific contacts with the various p53 response elements. The six amino acid residues that are most frequently mutated in human cancer are located in or close to the DNA-binding surface. These residues have been classified as ‘‘contact’’ (Arg-248 and Arg-273) or ‘‘structural’’ (Arg-175, Gly-245, Arg-249,andArg-282)residues, depending on whether they directly contact DNA or play a role in maintaining the structural integrity of the DNA-binding surface (Joerger, Ang, and Fersht 2006).

The DBD is structurally stabilized by Zinc atom as shown in figure 1, Zinc also acts as a metal chelator inducing the protein to adopt an immunological phenotype, and hence zinc is required for the proper folding of the protein. It was shown experimentally that the addition of extracellular zinc at a concentration within the physiological range (5µM) can cause renaturation and reactivation of wild-type p53 (Richard, Hainaut, and Méplan 2000).

**Figure 1:**
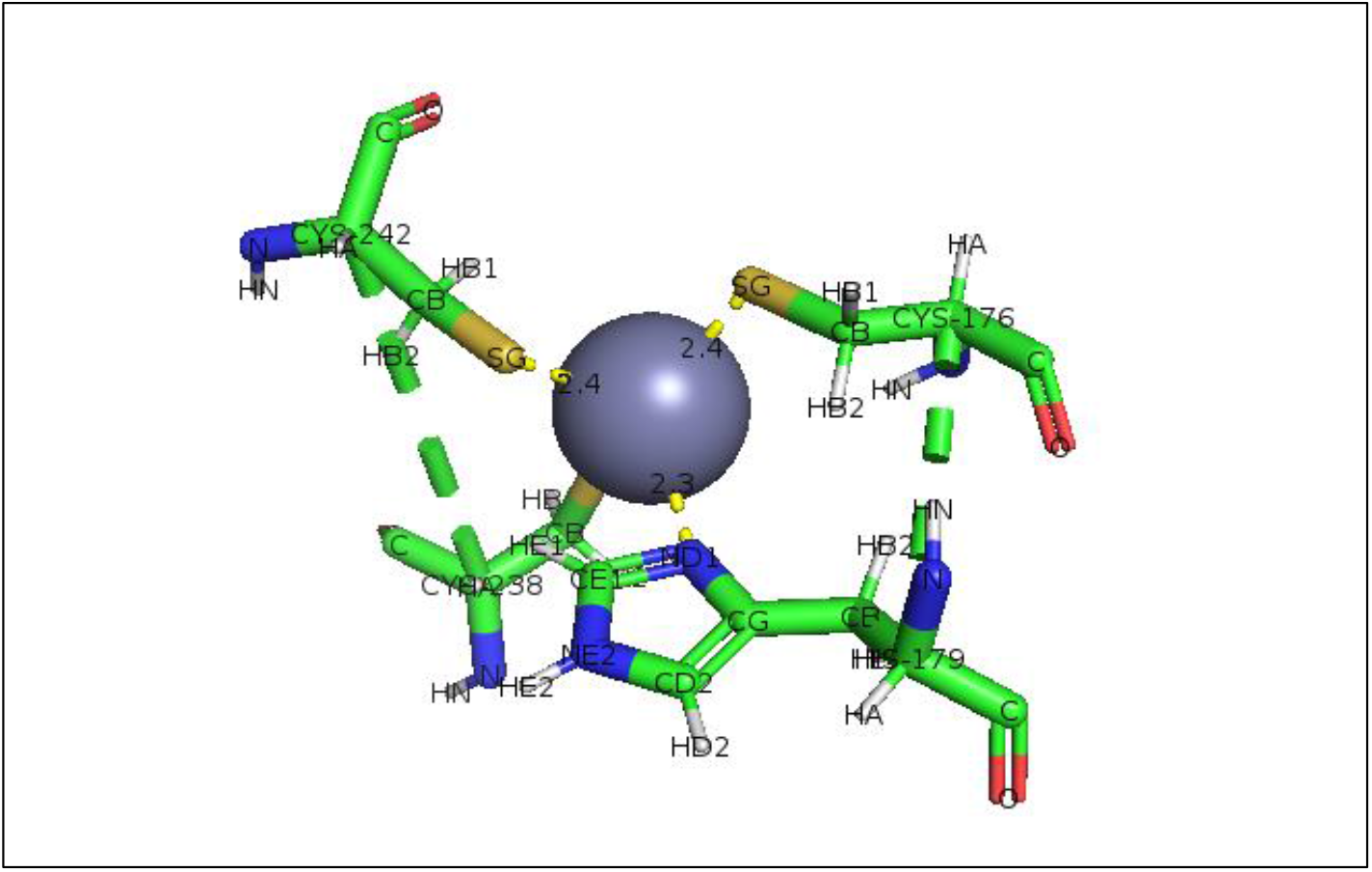
Zinc in p53 forms coordinate bond with 3 cysteine residues namely Cys176, Cys238, Cys242, and 1 His179 residue and structurally stabilizes its DNA binding domain (DBD).

Wild type p53 is predominantly in the holo (zinc-bound) form. The p53-Y220C mutant (our Target) is prone to the loss of Zn2+ due to local unfolding and increased aggregation, so the exact value of Kd is unknown (Miller et al. 2018).

Dimers are the predominant form of p53 in the cellular pool based on the estimated basal concentration of p53 in cells, Assembly of p53 into oligomers, represent a crucial factor for its tumor-suppressive function (Kitayner et al. 2006). It was experimentally determined that dimeric p53 variants are cytostatic and can arrest cell growth, but cannot trigger apoptosis in p53-null cells. In contrast, p53 tetramers induce rapid apoptosis and cell growth arrest, while a monomeric variant is functionally inactive, supporting cell growth. A shift in equilibrium between oligomeric states can be controlled through multiple known protein-protein interactions, either by the formation of tetramer or sequestering p53 in a monomeric state. p53 protein devoid of the tetramerization domain is mostly monomeric in solution but form tetramers with DNA targets incorporating 2 decameric repeats, Hence shifting the oligomeric state equilibrium of cell toward monomer or tetramer is a key parameter in p53-based cell fate decision (Fischer et al. 2016).p53 binds as tetramer to diverse DNA targets containing 2 decameric half-sites of general form RRRCWWGYYY(R=A, G; W=A, T; Y=C, T), separated by 0-13 base pairs In all structure, four p53 molecules self-assemble on 2 DNA half-site to form a tetramer that is dimer of dimers, Stabilized by protein-protein and base stacking interactions (McLure 2002).

In Y22OC mutant p53 tyrosine at 220^th^ residue is substituted by cysteine. It is located at the far end of the sandwich, at the start of the loop connecting strands S7 and S8 and destabilizes the core domain by 4kcal mol. The benzene moiety of Tyr-220 forms part of the hydrophobic core of the sandwich, whereas the hydroxyl group points toward the solvent. The 1.65-Å crystal structure of T-p53C-Y220C showed that the Y220C mutation creates a solvent-accessible cleft that is filled with water molecules at defined positions but leaves the overall structure of the core domain intact, The structural changes upon mutation link two rather shallow surface clefts, preexisting in the wild type, to form a long, extended crevice in Tp53-Y220C, which has its deepest point at the mutation site. Cys-220 occupies approximately the position of the equivalent atoms of Tyr-220 in the wild type, the mutation, however, results in a loss of hydrophobic interactions and a suboptimal packing of these hydrophobic core residues. The side chain of Leu-145, which was buried in the wild type, for instance, becomes partly solvent-accessible in Tp53-Y220C (figure 2). The largest structural changes in the immediate environment of the mutation site are found in the S7–S8 (Joerger, Ang, and Fersht 2006).

**Fig 2:**
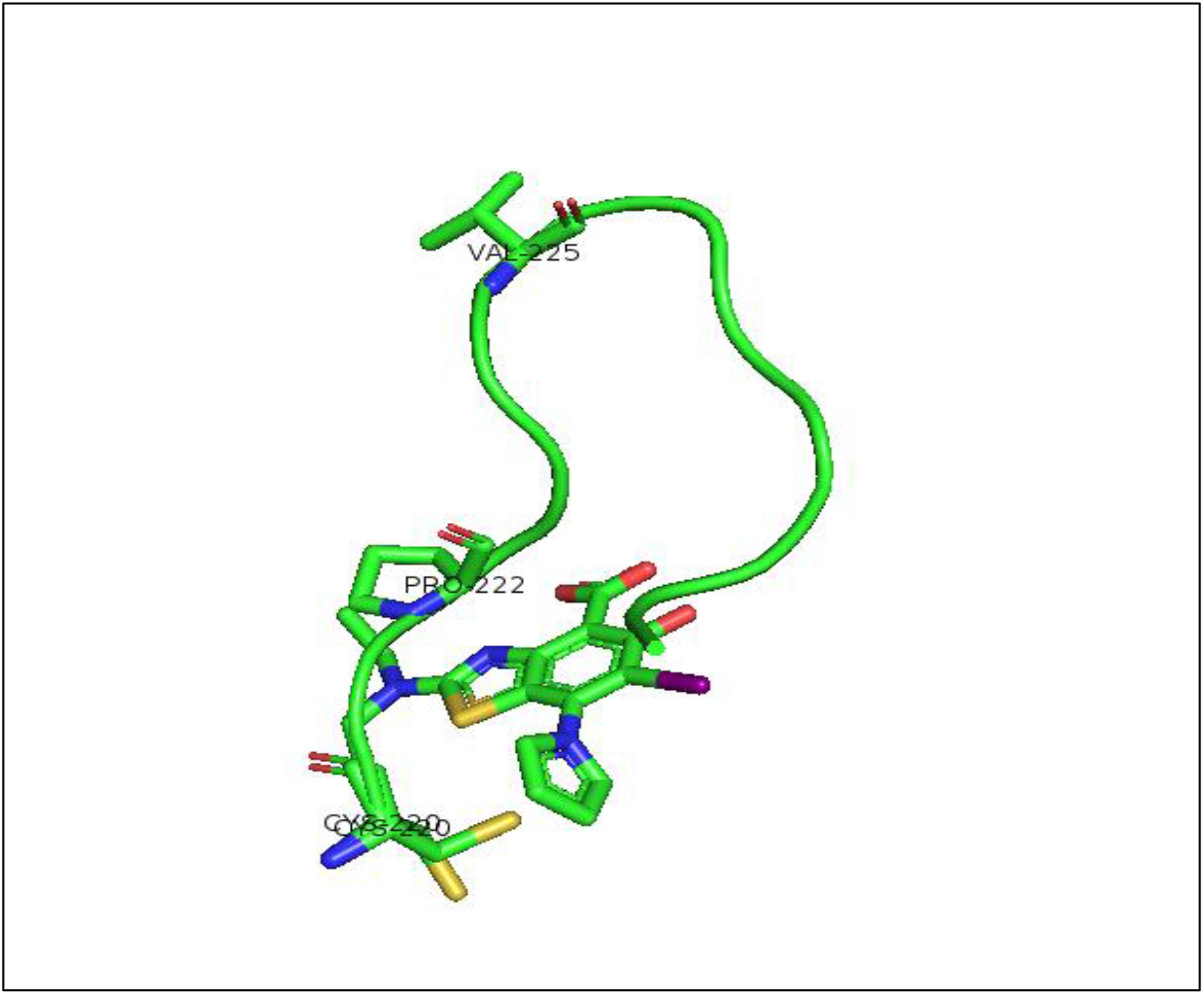
S7-S8 loop destabilized by mutation with stabilizer MB710 bounded to crevice formed by replacement of tyrosine by cystine. The maximum destabilization begins after Pro-222 residue exposing the hydrophobic core Val-225.

The binding site can formally be subdivided into a central cavity and 3 subsites 1, 2, 3. A narrow channel on the opposite end of the central cavity leads into subsite2, where several proteins including pro153, provide a hydrophobic interaction surface, with several backbone oxygens lining this hydrophobic patch (cys220, pro151, and pro152).Subsite3-At the bottom of the central cavity is essentially modulated by the conformational state of Cys220. various small molecules sample two different conformational states of the binding pocket that are essentially modulated by Cys220 Rotamer switch of the side chain of Cys220 from trans to gauche(-) conformation opens up a small hydrophobic pocket at the bottom of the binding site (subsite III). Conformational flexibility in the subsite III region is not restricted to Cys220 but also extends further into the hydrophobic core of the beta-sandwich region.Val157 and Ile232 adopt alternative side-chain conformations, depending on the bound ligand, which is indicative of the dynamic nature of this region of the binding pocket (Baud et al. 2015).

## 2. Material and Methods

### 2.1 Retrieval of protein structures

Protein structures of the DBD of wild type p53 with PDB ID **2fej** and mutant p53 bound to ligand with PDB ID **501i** were retrieved from Protein Data Bank (http://www.rcsb.org).

### 2.2 Molecular Dynamic Simulation studies of wild type p53, mutant p53, and mutant bound to Ligand

Protein structures were retrieved from the RCSB PDB database and missing atoms, residues and loops were completed i.e. protein was prepared using Discovery Studio 3.1 prepare protein option. Protein was simulated using the gromos54a7 force field (http://www.gromacs.org/). Ligand topology was generated using the PRODRG server. The prepared protein-ligand complex was solvated in a dodecahedron box with a box length of 1.0nm using the SPC216 water model. The overall system was neutralized by adding 7 chlorine in the case of wild type p53, 4 chlorine in case of mutant p53 and 5 chlorine for mutant p53 bound to the ligand as counter charge ions and relaxed with 50000 steps of steepest descent minimization. Equilibration of the system was performed in two ensembles, NVT (constant number of volume, temperature & pressure) ensemble which ran for 100 picoseconds stabilizes the temperature of the system and NPT (constant number of particles, pressure & temperature) ensemble which ran for 100 picoseconds stabilizes the pressure & density of the system. Upon completion of two equilibration phases, finally, we ran production MD for 10 nanoseconds & results were analyzed.

### 2.3 Distance restraining the zinc atom

p53 protein has a zinc atom in its DNA binding domain, It was seen that zinc atom was lost at the end of simulation and as zinc atom is crucial for proper functioning of p53, zinc atom was restrained, for retraining zinc atom coordinate bond between zinc and three cysteines and one histidine was measured, also the atom and residue number and specific interactions were noted and using this information a table was made.

### Using this information the Distance restraint matrix was generated as followed…

|  |  |  |  |  |  |  |  |  |
| --- | --- | --- | --- | --- | --- | --- | --- | --- |
| ; ai | aj | type | index | type' | low | up1 | up2 | fac |
| 3169 | 1328 | 1 | 0 | 1 | 0.0 | 0.22 | 0.24 | 1.0 |
| 3169 | 1286 | 1 | 1 | 1 | 0.0 | 0.23 | 0.25 | 1.0 |
| 3169 | 2249 | 1 | 2 | 1 | 0.0 | 0.23 | 0.25 | 1.0 |
| 3169 | 2295 | 1 | 3 | 1 | 0.0 | 0.23 | 0.25 | 1.0 |

This matrix was added in the topology file and a similar process was repeated for mutant as well as for mutant bound to the ligand.

### 2.4 Retrieval of known Inhibitors for the Y220C mutant target and Molecular Docking

All the known small molecules were retrieved from the literature. Structures were drawn by Marvin’s sketch (https://www.chemaxon.com/marvin). Further, these drawn structures were used for docking, based on available data of activity five protein structure were chosen and all the compounds were docked into them, the chosen protein was of 2 categories open type and closed type, Docking was carried out using Lib dock software of Discovery studio 3.1

### 2.5 Calculation of Descriptors

Descriptors are numerical values that characterize the property of molecules. The data set should contain at least 5 times as many Compounds as descriptors in QSAR; the reason being too few compounds relative to the number of descriptors will give a falsely high correlation

Descriptors fall into four classes

a. Topological
b. Geometrical
c. Electronic
d. Hybrid or 3D descriptors

Topological Descriptors are derived directly from the connection table representation of the structure which includes Atom and Bond Counts, Substructure Counts, etc. We used only topological descriptors for our work.

Based on Dock score and activity data, 5 small molecules were chosen for descriptor calculations. Descriptor was calculated using Pharmaceutical Data exploration laboratory PADEL software (http://www.yapcwsoft.com/dd/padeldescriptor/), Descriptors chosen were Molecular weight, number of aromatic rings, number of atoms, number of heavy atoms, number of rotatable bonds, no. of iodine, presence or absence of pyrrole.

### 2.6 Virtual Screening of Drug bank Database

The descriptors were used to screen the complete Drug bank database. Drug bank had about 9300 compounds and descriptors were used to separate the database into actives and in actives, about 5000 compounds were separated into actives. All the 5000 compounds were docked into the same target (5o1i) and Ranked using Discovery studio 3.1 and compounds getting a lib dock score of minimum 135 or above were used for further analysis. Top 5 compounds were selected; 2D & 3D interaction maps were generated using Discovery studio and Pymol (https://pymol.org/2/).

### 2.7 Molecular Dynamic Simulation studies of Top Compound from Virtual screening output

Output from virtual screening were undergone 2D and 3D interaction studies to check specific interactions also their literature studies were carried out to understand their targets and mode of action and on the basis of this and dock scores, 2 compounds were made into complexes with mutant p53 protein (PDB ID 501i) using pymol software and complexes were simulated using gromos54a7 force field. Ligand topology was generated using the PRODRG server. The prepared protein-ligand complex was solvated in a dodecahedron box with a box length of 1.0nm using the SPC216 water model. The overall system was neutralized by adding 6 chlorine for 1st compound and 7 chlorine in case of 2^nd^ compound as counter charge ions and relaxed with 50000 steps of steepest descent minimization. Equilibration of the system was performed in two ensembles, NVT (constant number of volume, temperature & pressure) ensemble which ran for 100 picoseconds stabilizes the temperature of the system and NPT (constant number of particles, pressure & temperature) ensemble which ran for 100 picoseconds stabilizes the pressure & density of the system. Upon completion of two equilibration phases, finally, we ran production MD for 10 nanoseconds & results were analyzed.

### 2.8 Fragment-based optimization and validation

The drug was optimized by overlapping with the most potent binders and then adding pharmacophores to maintain important interactions, again the analogs were docked back into the same target and simulated and compared to results of unmodified drugs and potent binders to validate the increase in activity also the analogs were submitted to SwissADME (http://www.swissadme.ch/) Server to predict ADME parameters, pharmacokinetic properties, druglike nature, and medicinal chemistry friendliness.

## 3 Results and Discussion

### 3.1 Molecular Dynamic Simulation

To investigate the structural dynamics of wild type p53 in comparison with the Y220C mutant protein we employed molecular dynamics simulations of the DNA binding domain from available X-ray data. Our modeling result predicts that Zinc atom was successfully distant restrained in case of wild type p53 but couldn’t be restrained in case Y220C mutant p53. Simulation curve shows that Wild type p53 shows instability in its unbound form but gets highly stabilized when bound to DNA (figure 3) which is in accordance to research paper which advocates that p53 monomer is in transient state and its tetramerization state decide the cell fate, p53 stabilizes only after binding to its specific response element.

**Fig 3:**
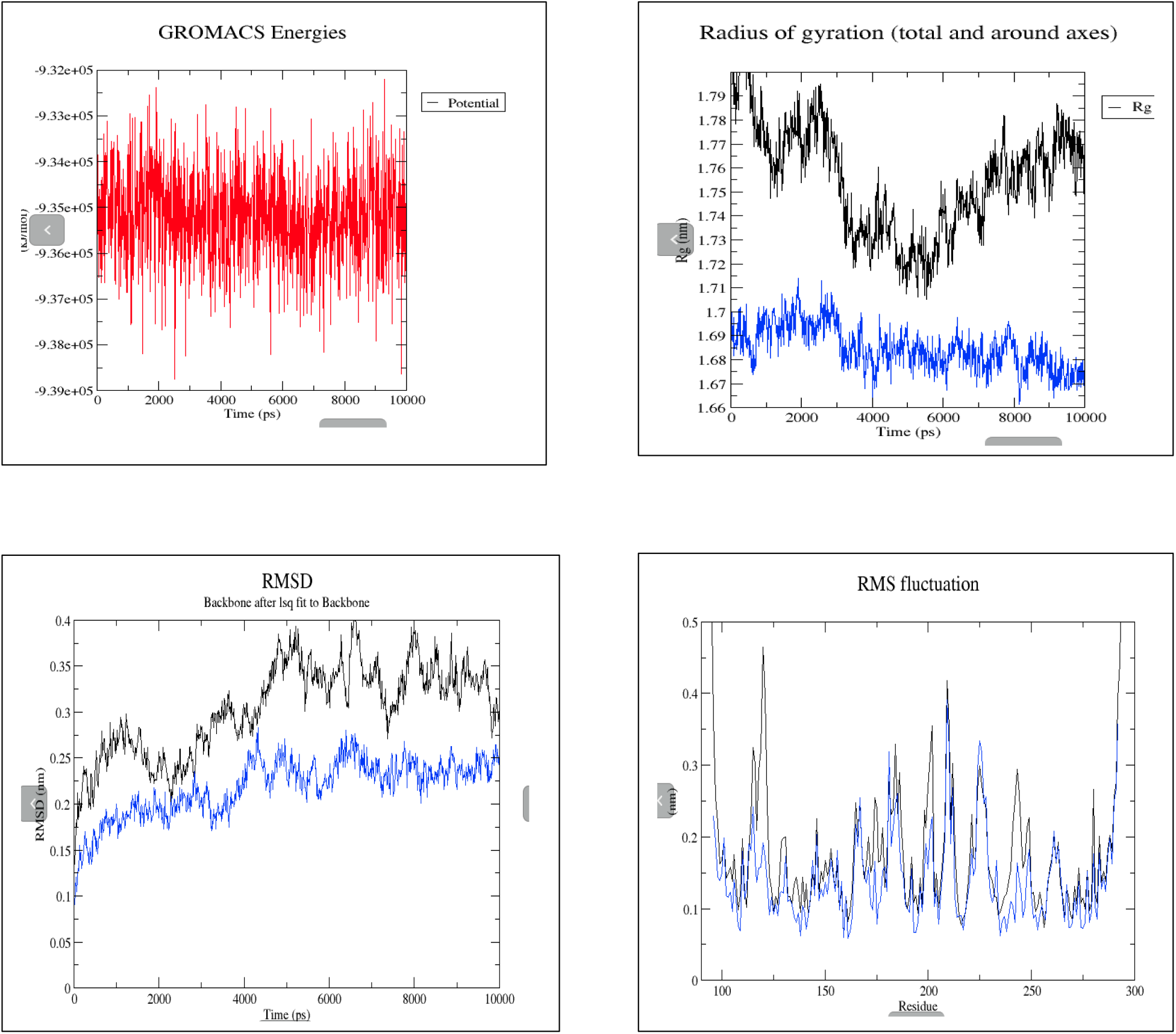
Simulation comparisons between wild type p53 (Black) and wild type p53 (blue) bound to DNA

GYRATION, RMSD and RMSF curves of Y220C mutant P53 bound to stabilizer MB710 shows to stabilize the mutant protein (figure 4). Mutation takes place at residue 220 where tyrosine is replaced by cysteine but RMSF curve shows fluctuation starting from residue 222 and it reaches its peak at residue 225, which is in accordance to research paper which advocates that mutation at residue 220 destabilizes the complete S7-S8 loop with maximum structural effect starting from residue 222 which is proline and the peak reaches its maxima at residue 225 which is a valine, which was buried in the hydrophobic core protected by the aromatic ring of tyrosine but Y220C mutation exposes the hydrophobic residue to the hydrophilic exterior.

**Fig 4:**
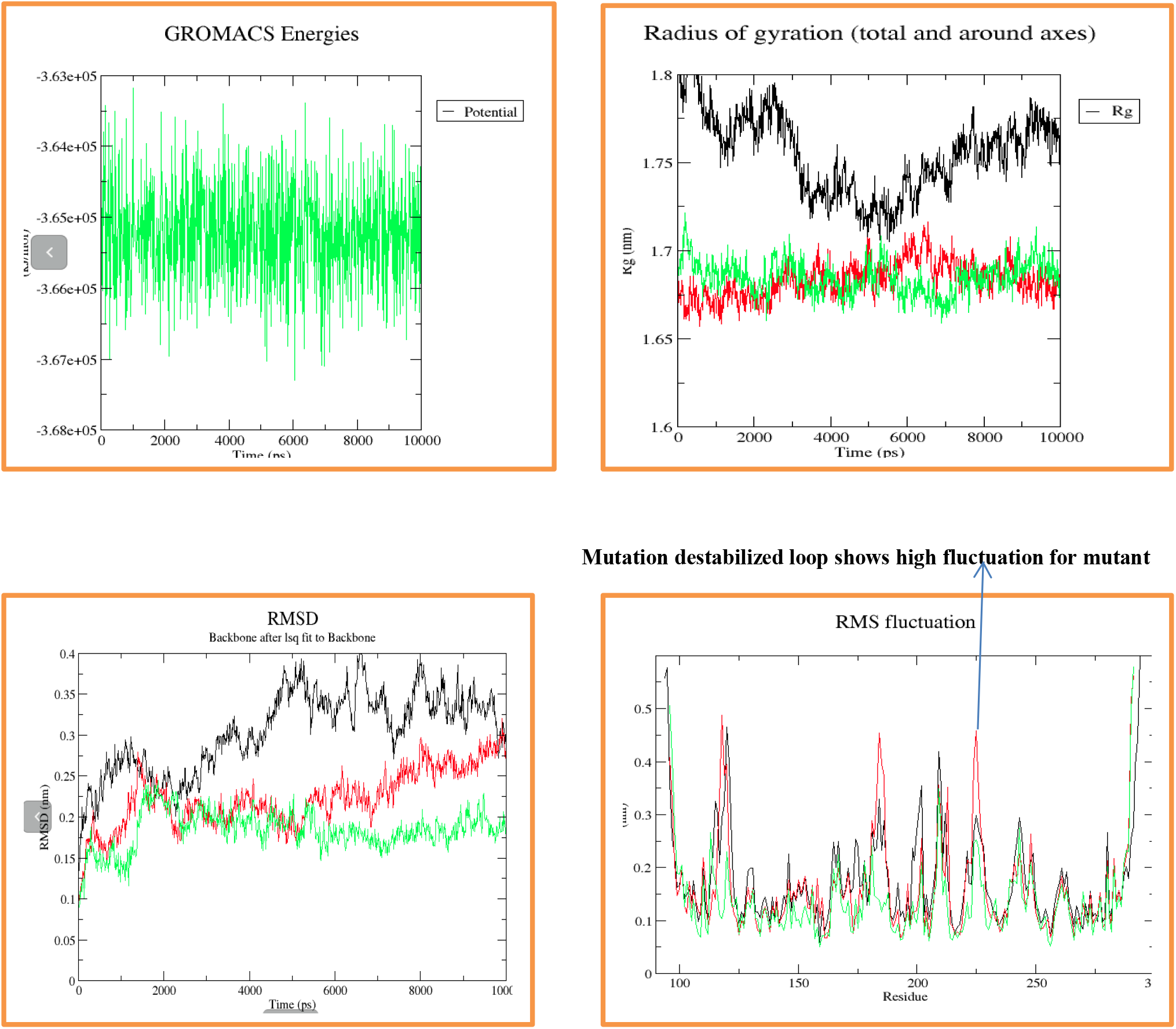
Simulation comparisons between wild type p53 (Black) with mutant (Red) and mutant bound to ligand MB710 (green)

## 3.2 Molecular Docking

To investigate the potential binding mode of all known ligands from literature studies with the mutation-induced cavity in Y220C mutant p53 we employed molecular docking, data given in Table 3. From the SAR library, 5 compounds with PDB IDs 501i, 4Agq, 2Vuk, 3ZME & 501G were given special focus based on literature study, in vitro activity and Dock score.

### 3.3 Virtual Screening Output

Drug Bank had around 9300 compounds and calculated descriptors (Table 2) were used for Virtual Screening and output of Virtual Screening was around 5000 compounds which were docked back into the same target and a cut off score of 135 on the basis of previous data was used as a threshold and 5 compounds were selected (Table 4). Among the compounds Masoprocol was given special attention because of its relative simpler structure which makes substitution easier also according to previous studies Masoprocol was found to induce apoptosis in tumor xenograft although its target and mode of action is unknown but it is well tolerated in animals and hence can be used for selective targeting of cancer cells without affecting the normal cells also masoprocol have structural similarities with small molecule analog L4 and L5 which were also designed against mutant p53 (Miller et al. 2018).

**Table 1:**
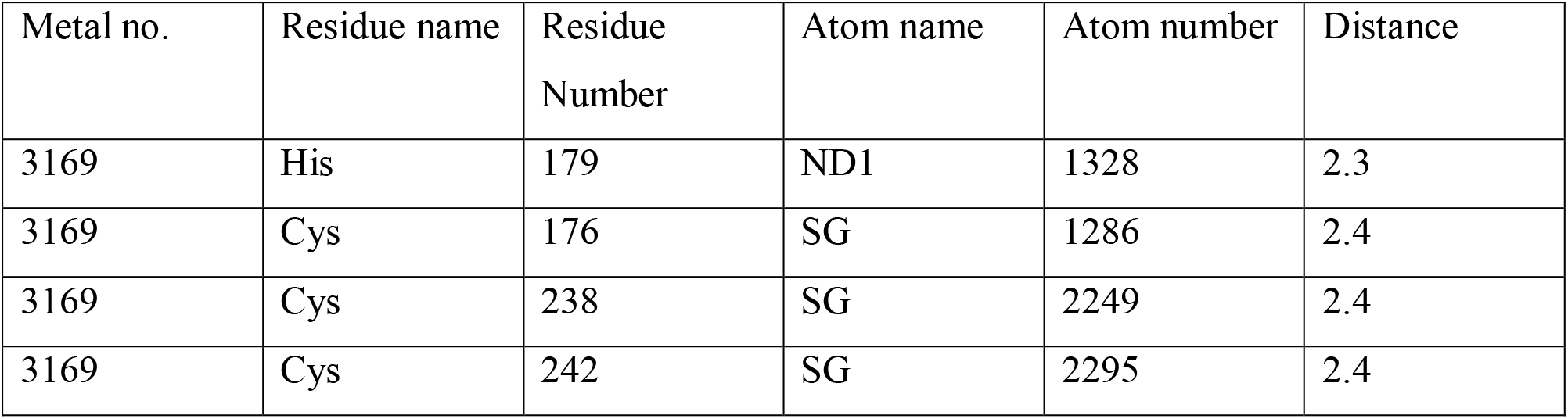
Distant Restraint parameters used to the restraint Zinc atom

| Metal no. | Residue name | Residue Number | Atom name | Atom number | Distance |
| --- | --- | --- | --- | --- | --- |
| 3169 | His | 179 | ND1 | 1328 | 2.3 |
| 3169 | Cys | 176 | SG | 1286 | 2.4 |
| 3169 | Cys | 238 | SG | 2249 | 2.4 |
| 3169 | Cys | 242 | SG | 2295 | 2.4 |

**Table 2:** Descriptors used to screen the Drug Bank Database

| Compound | Molecular weight | N-ArR | n-Atoms | Num-RotB | nI |
| --- | --- | --- | --- | --- | --- |
| 501i(open) | 457.290 | 3 | 40 | 5 | 1 |
| 3ZME(open) | 298.365 | 3 | 41 | 5 | 0 |
| 501g(open) | 449.244 | 3 | 41 | 6 | 1 |
| 2Vuk(close) | 238.334 | 3 | 36 | 3 | 0 |
| 4Agq(close) | 517.455 | 3 | 62 | 7 | 1 |

**Table 3:**
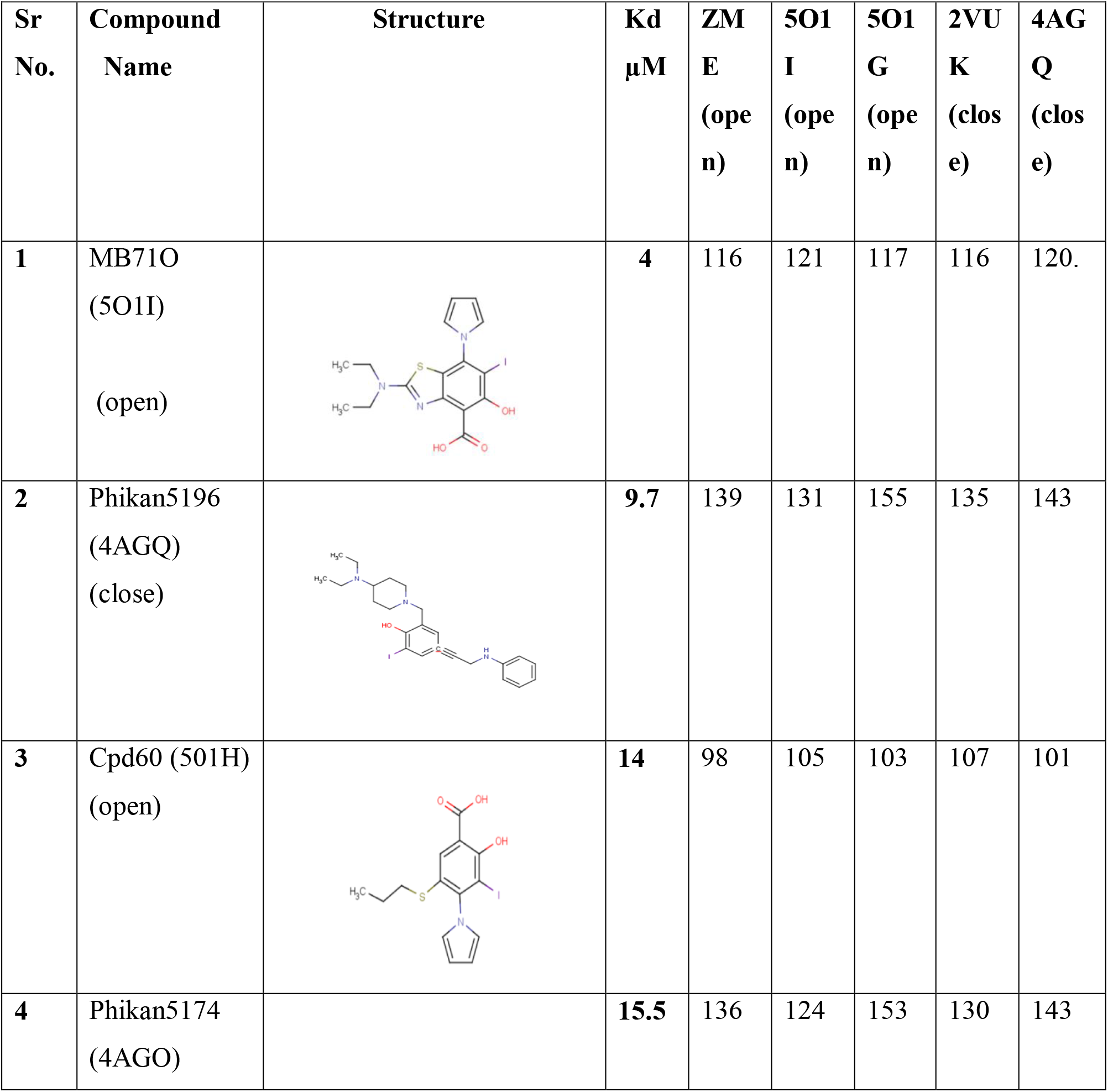
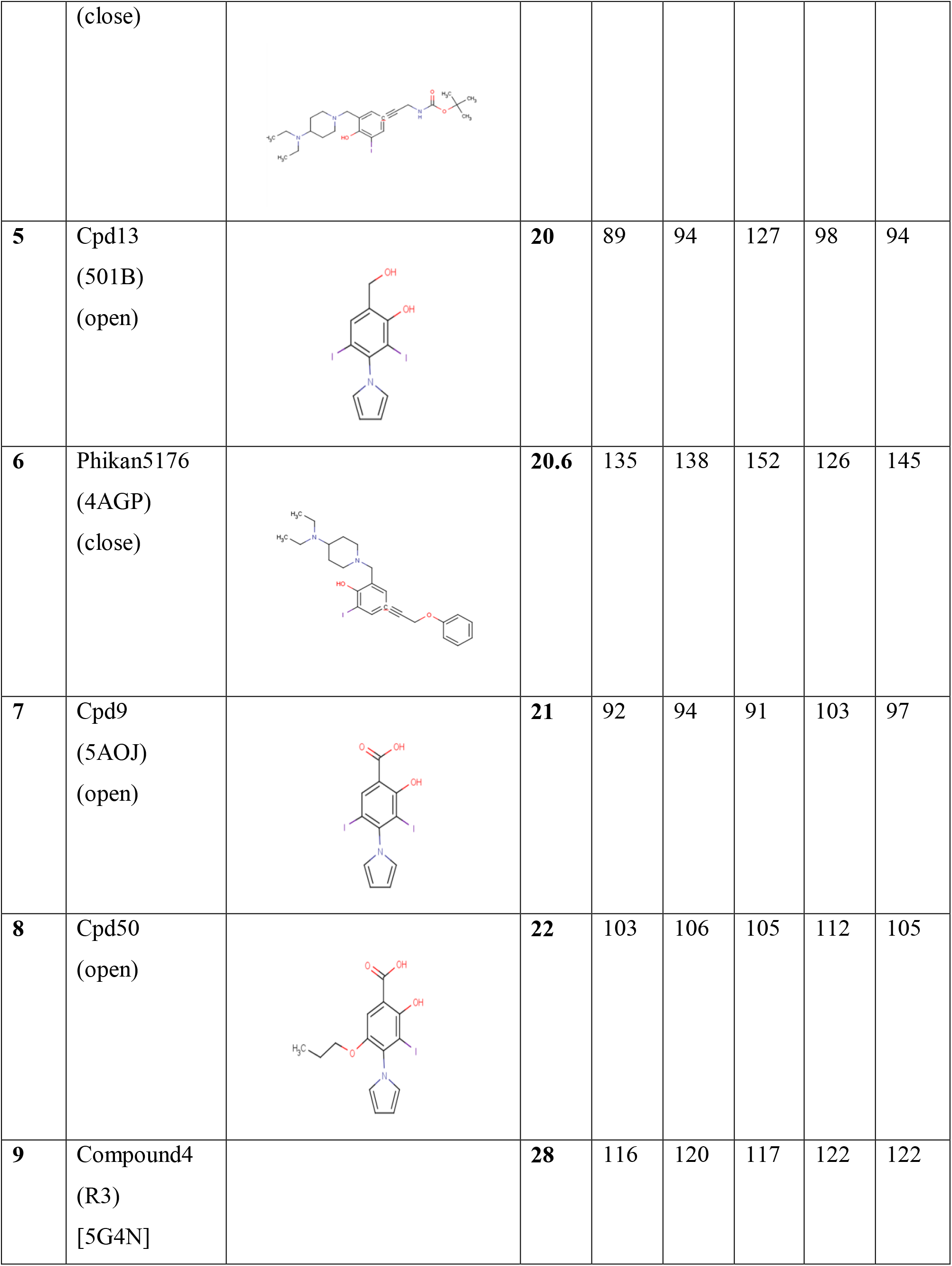
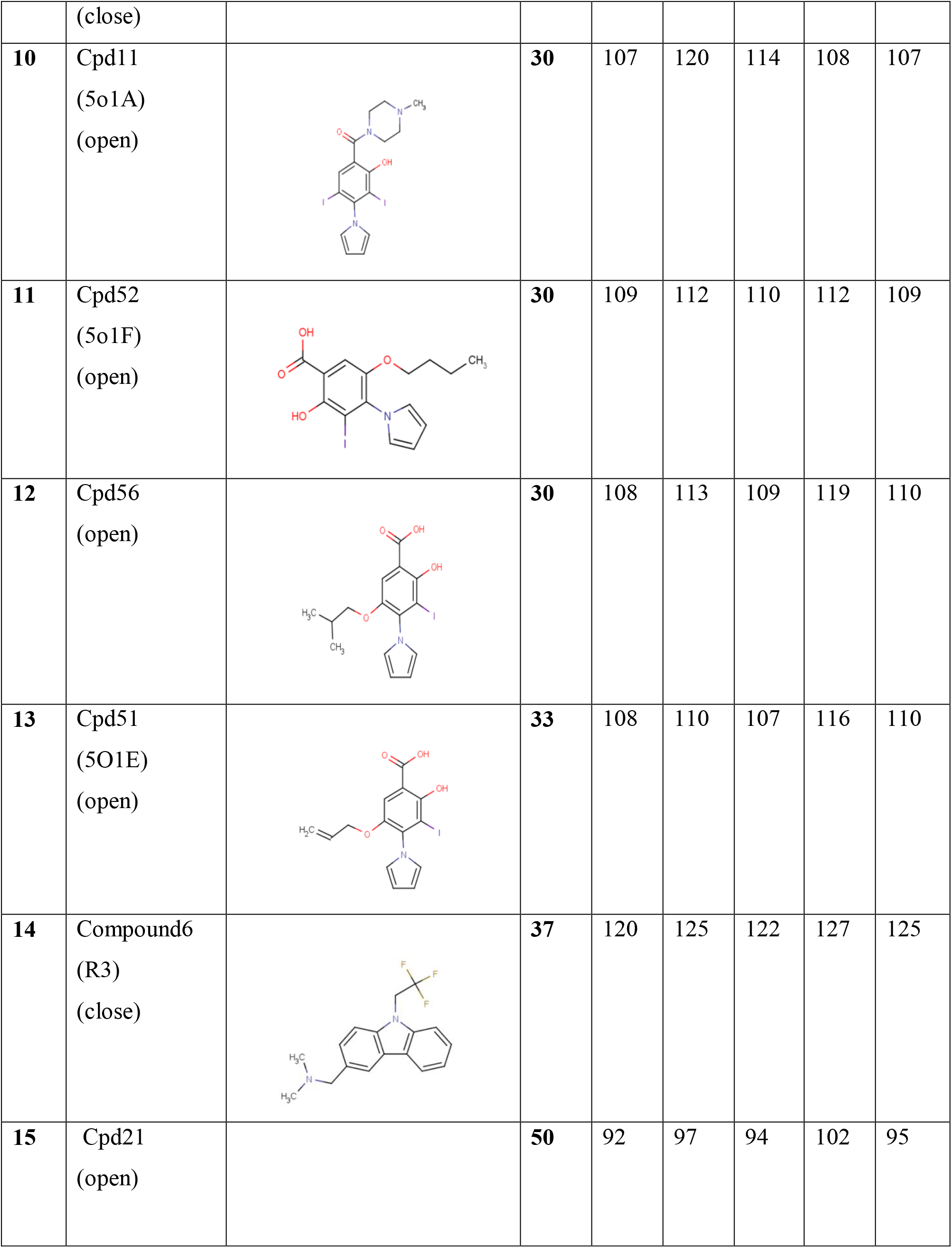
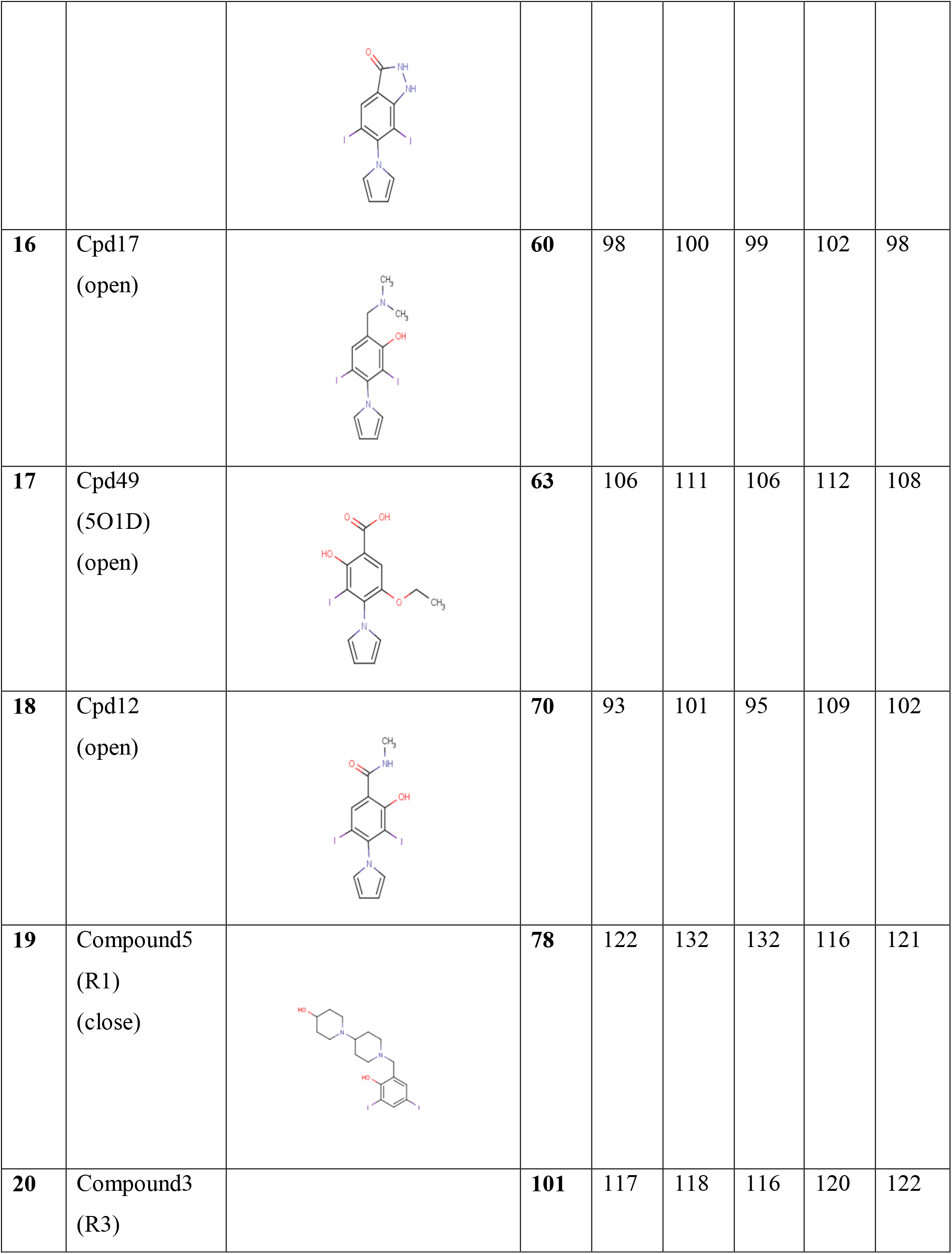
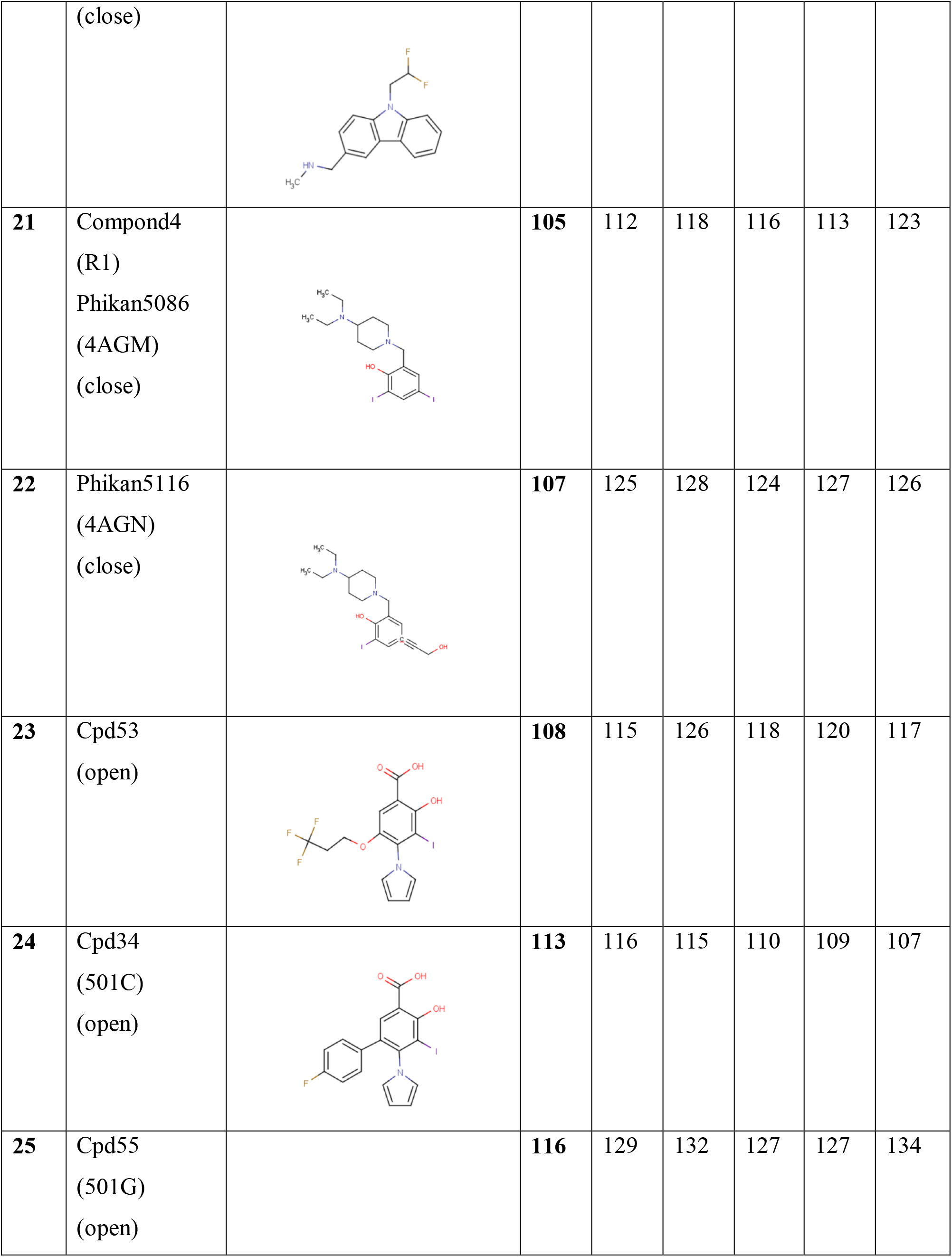
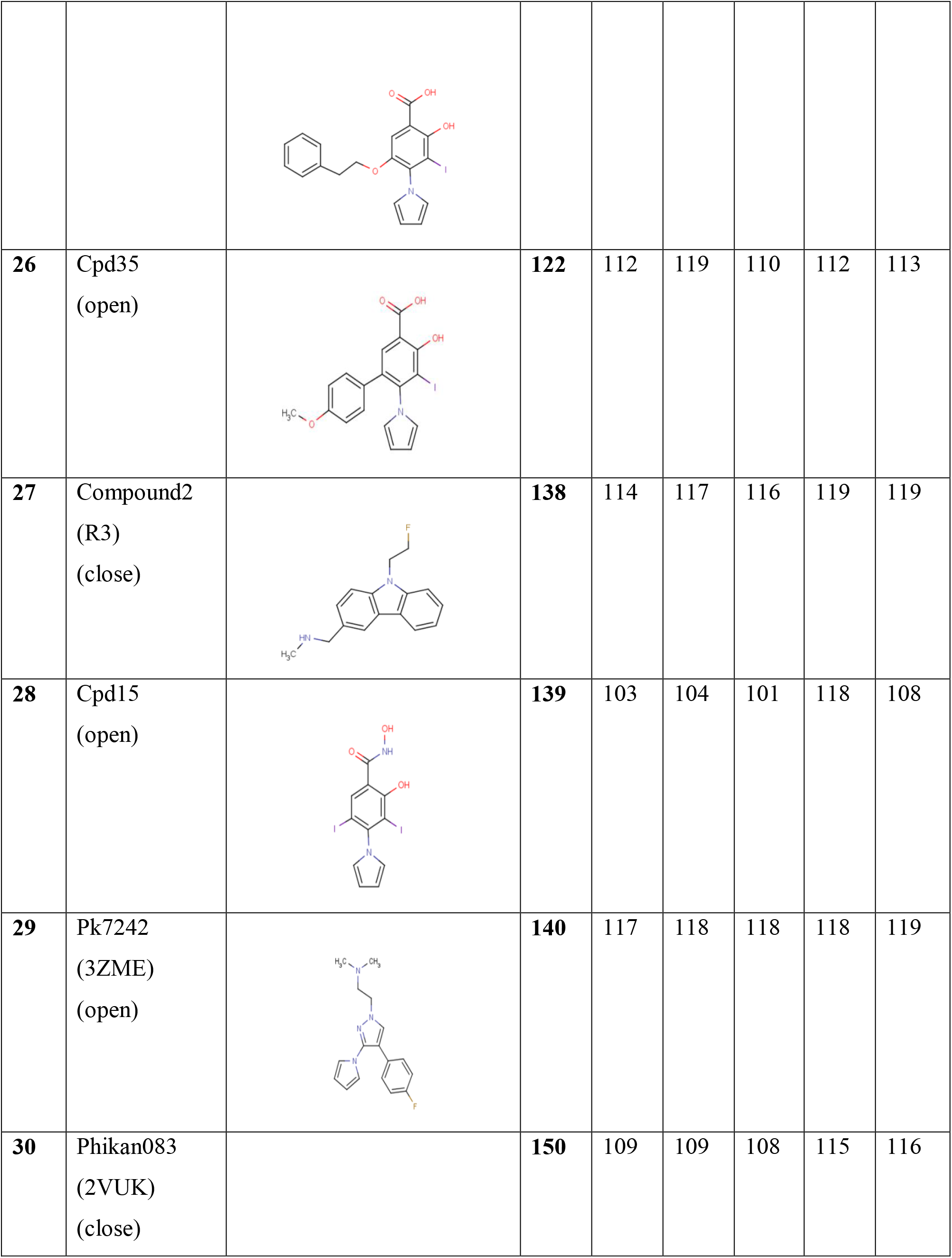
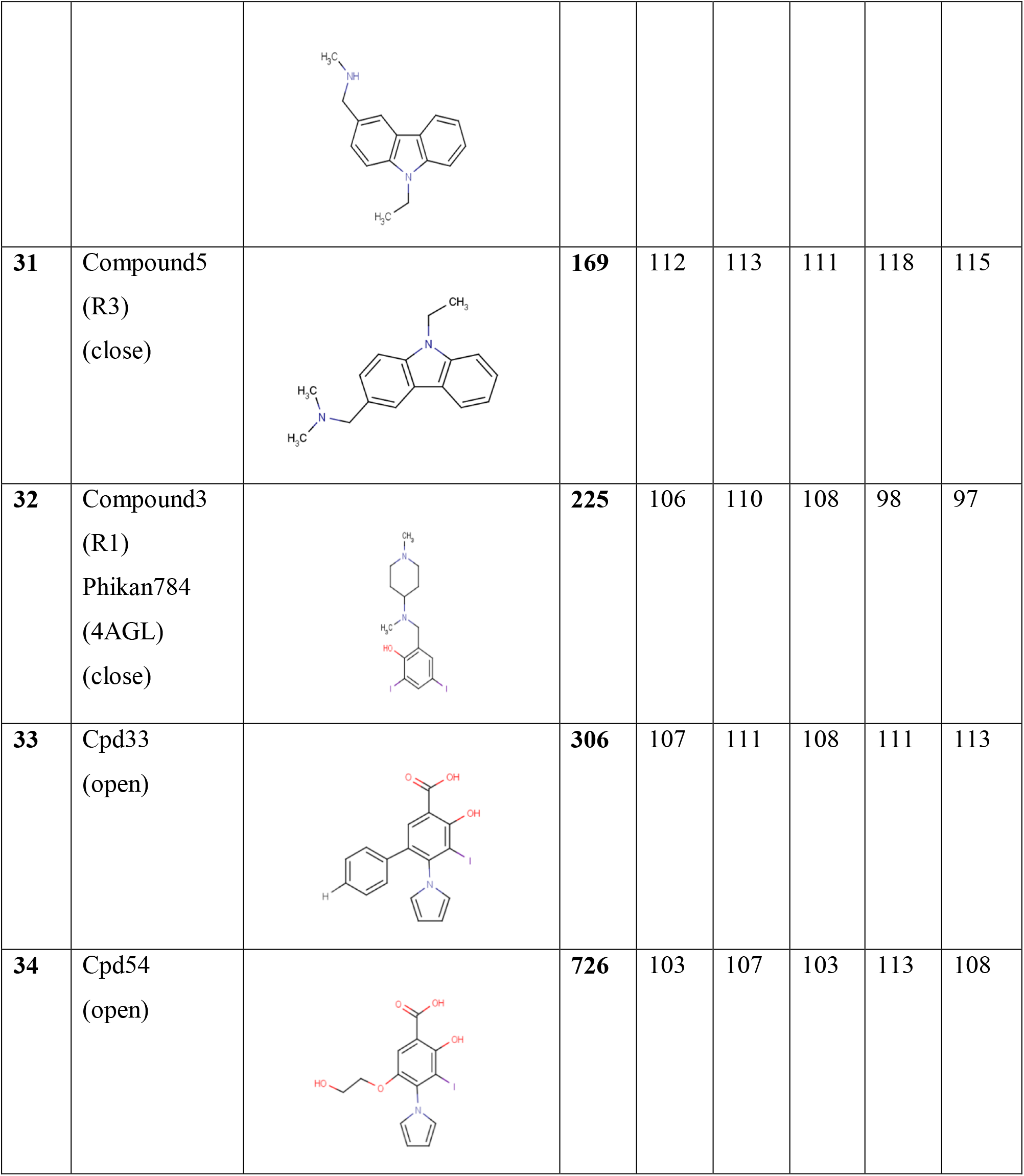
Based on Dock score and activity values as retrieved from literature ligand library was made from higher to lower activity

**Table 4:**
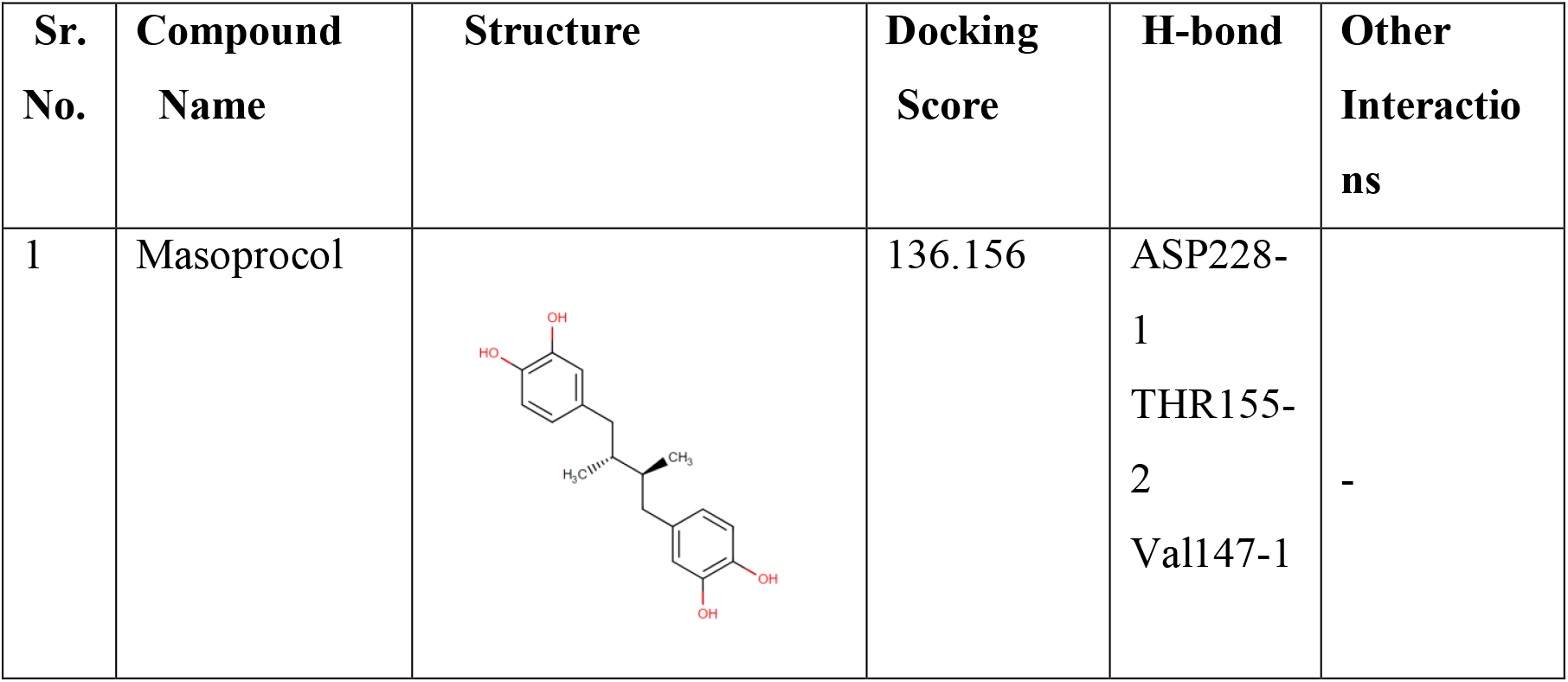
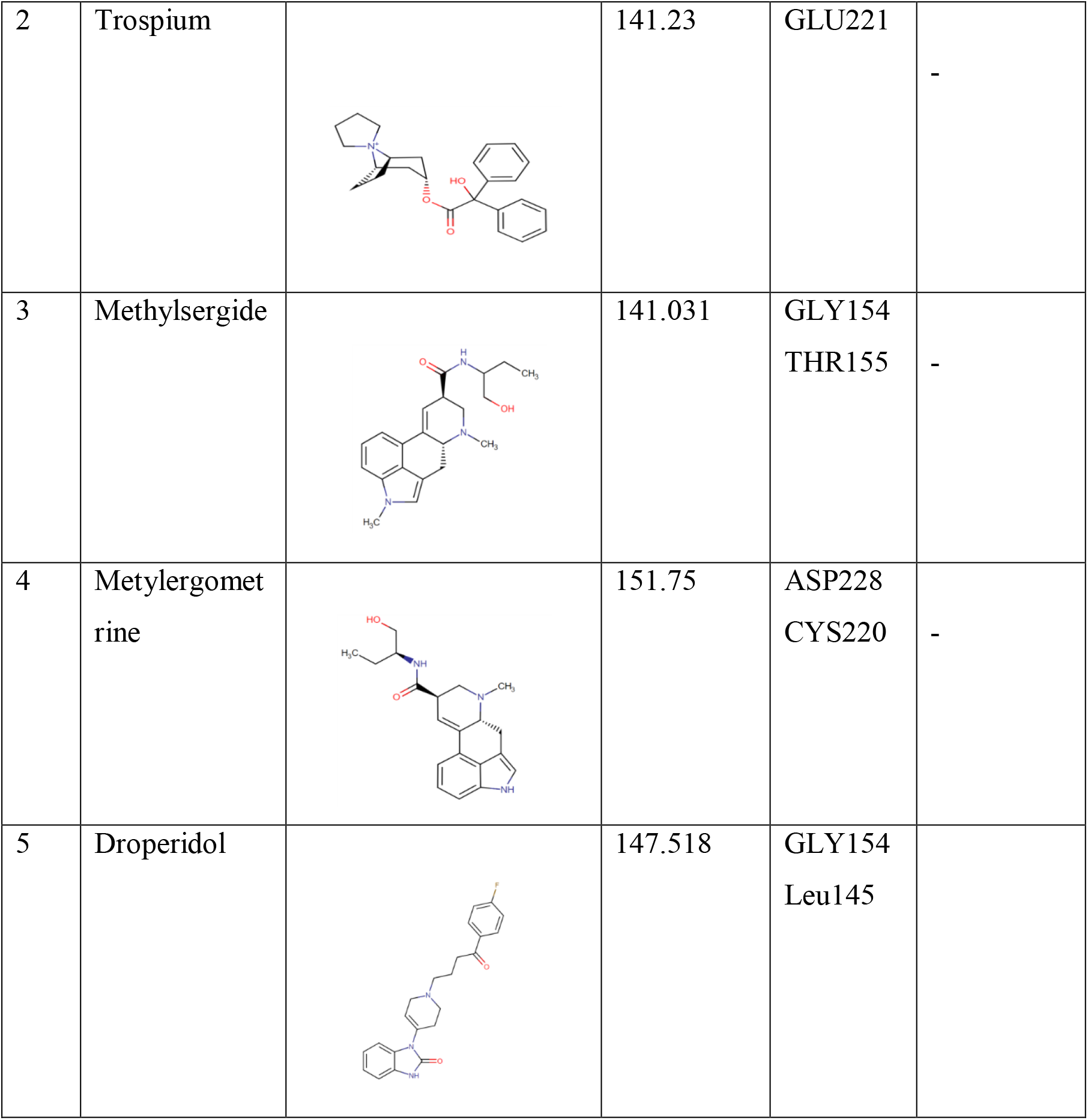
Top compounds out from Virtual Screening

| Sr. No. | Compound Name | Structure | Docking Score | H-bond | Other Interactions |
| --- | --- | --- | --- | --- | --- |
| 1 | Masoprocol |  | 136.156 | ASP228-1<br>THR155-2<br>Val147-1 | - |
| 2 | Trospium |  | 141.23 | GLU221 | - |
| 3 | Methylsergide |  | 141.031 | GLY154<br>THR155 | - |
| 4 | Metylergometrine |  | 151.75 | ASP228<br>CYS220 | - |
| 5 | Droperidol |  | 147.518 | GLY154<br>Leu145 |  |

Masoprocol got a Dock Score 136.156 which was higher than the Dock score of reference compound (501i) 121.355 But simulation results of Masoprocol vs. 501i showed that Masoprocol gyration was very similar to reference compound but its RMSD and RMSF curves showed much higher deviation and fluctuation (fig 6) clearly indicating that Masoprocol interactions with the binding site residues are not stable and Masoprocol needed structural optimization

### 3.4 Interaction Studies of Masoprocol

2D and 3D interaction maps were generated for output compounds from virtual screening (figure 5) and studied to see whether specific interactions are there or missing, based on interaction and literature studies Masoprocol was selected and analyzed

**fig 5:**
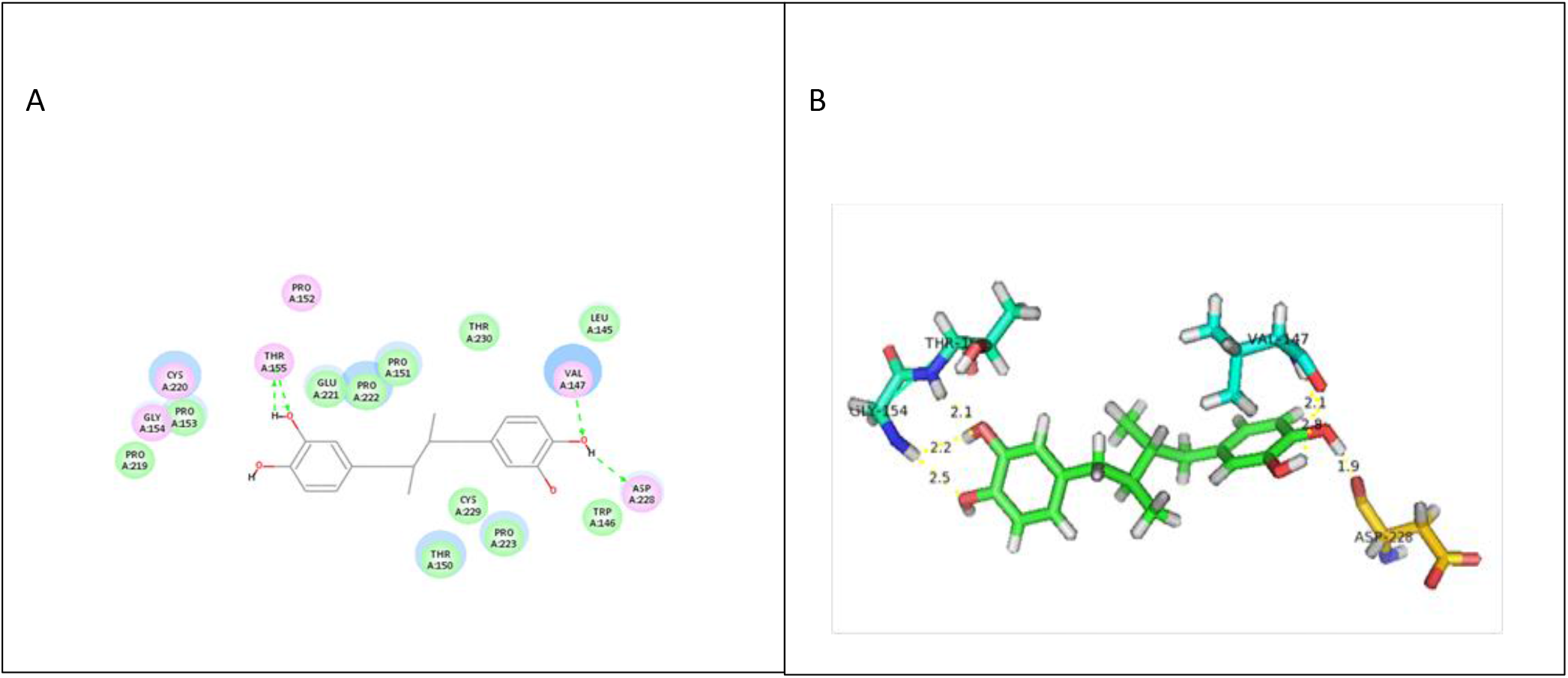
2D & 3D interaction map of masoprocol with the target (501i)

**Fig 6:**
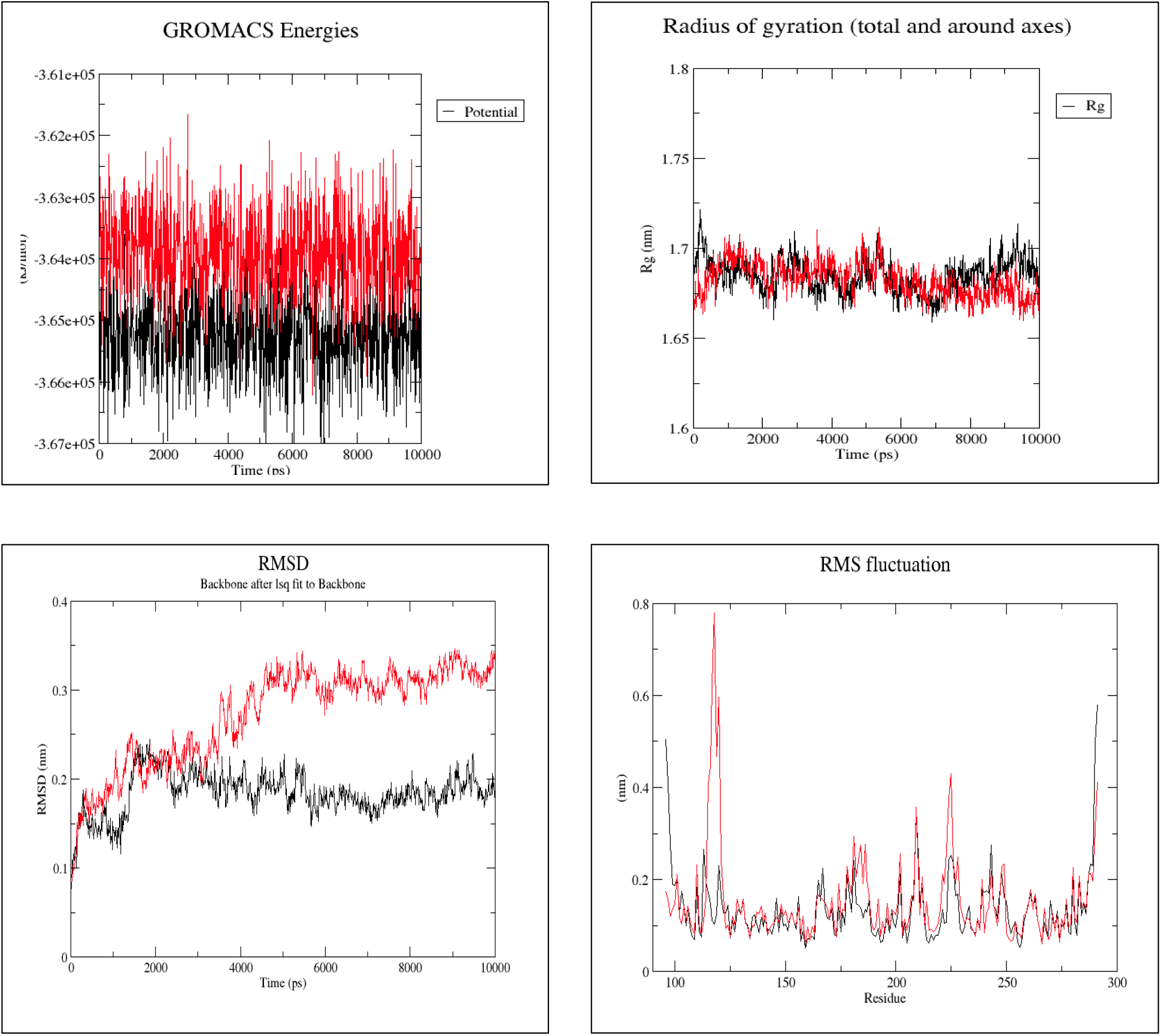
Simulation comparisons between mutant p53 bound to Masoprocol (RED) and mutant p53 bound to MB710 (BLACK)

### 3.5 Molecular Dynamic simulations of masoprocol in comparison with mutant p53 bound to MB710

## 4. Fragment-Based Optimization of Masoprocol

It is evident from the results of the simulation of Masoprocol Vs. MB710 that interactions of masoprocol with the binding site wasn’t stable and needed further optimization, for optimization 2 different schemes were followed, in scheme 1 (figure 7) pyrrole scaffold was added specifically by overlapping with reference ligand MB710.

**Fig 7:**
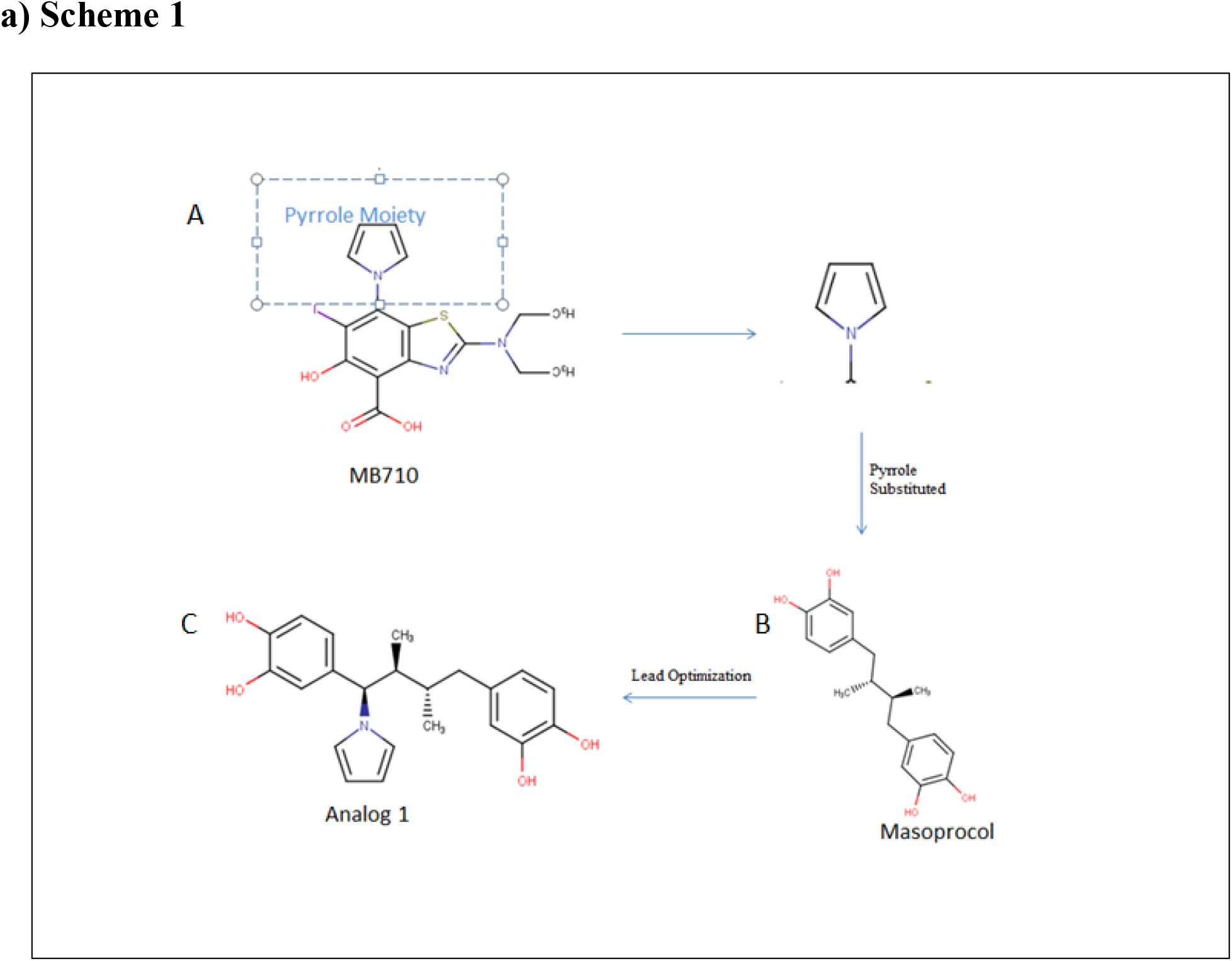
A) MB710 is the strongest binder B) Masoprocol found as a result of V.S C) Substituted Masoprocol-Analog1

In scheme 2 (figure 8) the benzothiazole N-ethyl scaffold was added to analog 1 to give rise to analog 2, again the groups were added after overlapping with reference molecule to maintain specific interactions. Benzothiazole scaffold connects the central cavity and subsite 2 via a conformationally restricted, aromatic sulfur heterocycle and the N-ethyl group targets a hydrophobic hotspot in subsite 2 formed by Pro151, Pro152, Pro153, and Thr155.

**Fig 8:**
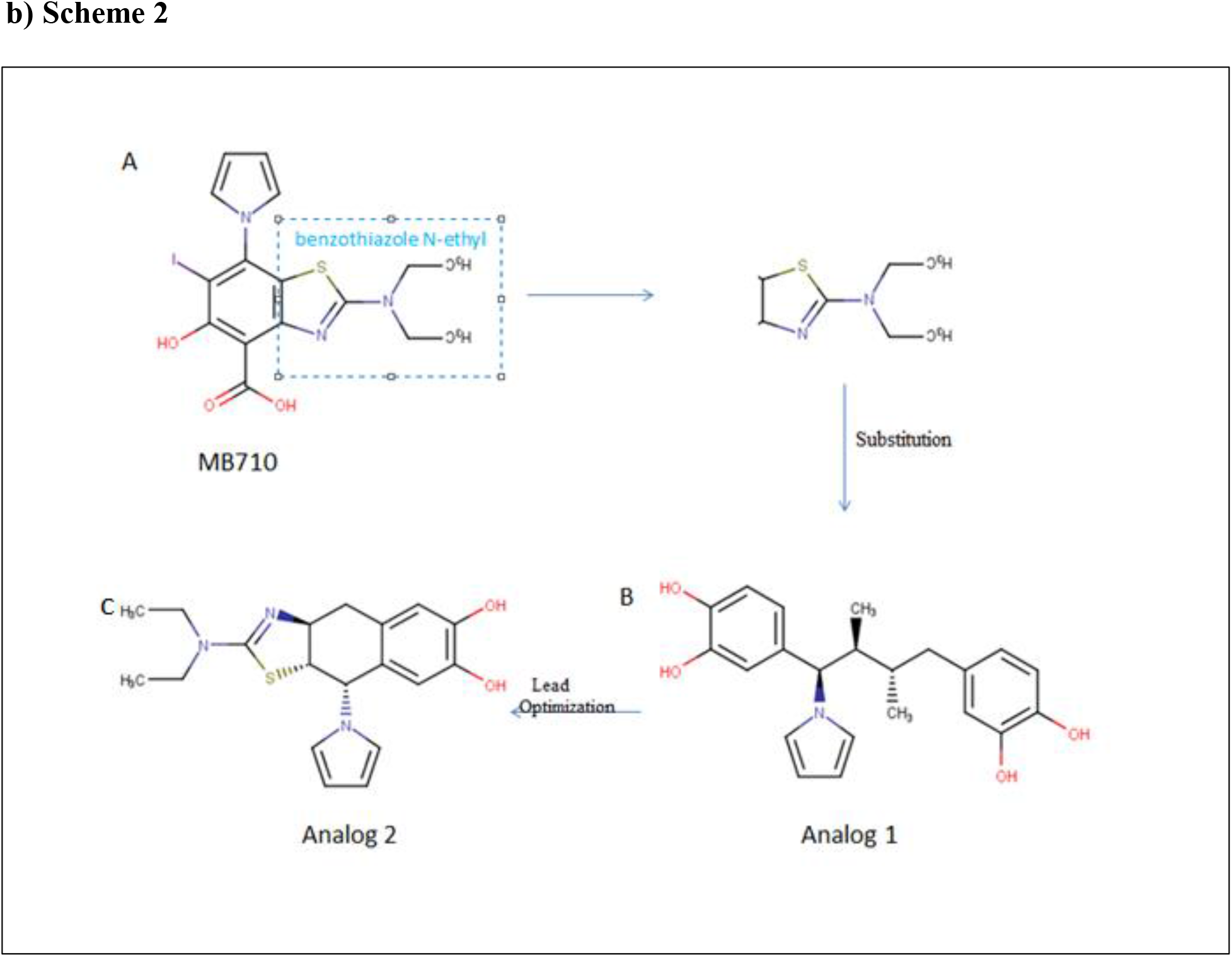
A) Benzithiazole group from MB710 and N-ethyl group also common in phikan5196 B) Substitution of pyrrole and benzothiazole N-ethyl scaffold into Analog1 C) Substituted Analog2

### 4.1 Validation of Masoprocol Analogs

The Analogs were again Docked against the same target, formed into complexes and simulated and simulation results were compared to Masoprocol (parent compound) and MB710 (present strongest binder i.e. reference Compound) to access their efficiency

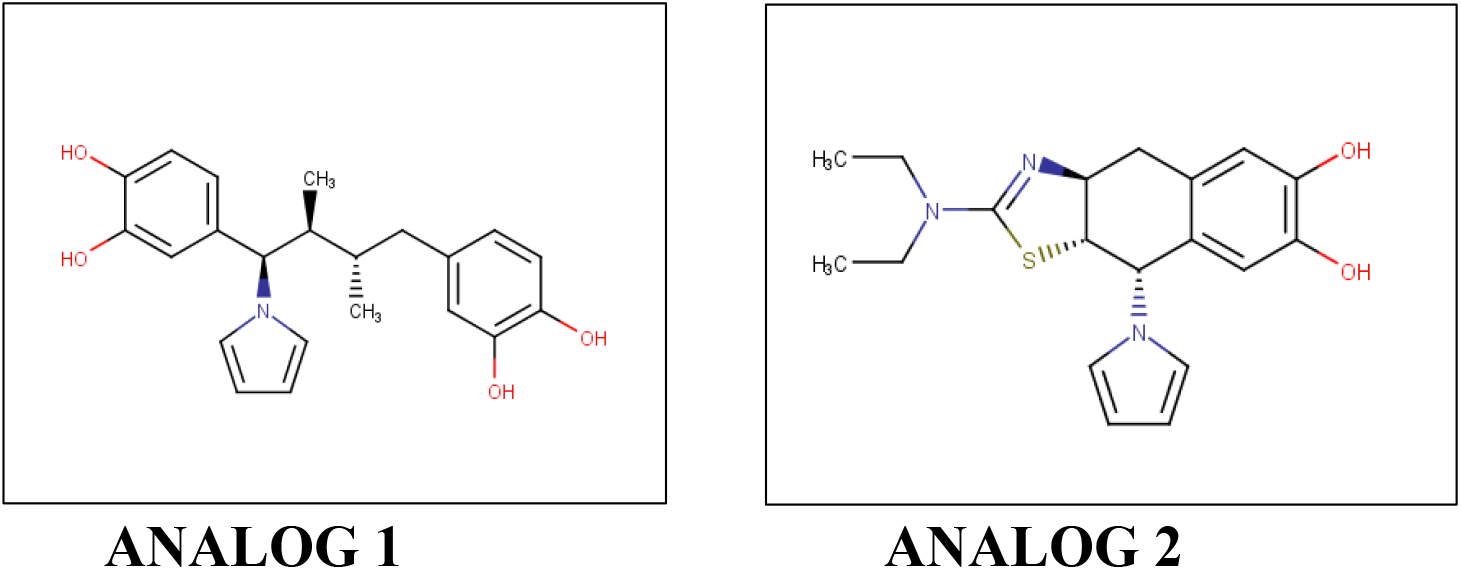

### 4.2 Interaction studies of Masoprocol Analogs

2D and 3D interaction maps were generated (figure 9) for Analog 1 and 2 to note down whether the important interactions are being maintained or not

**fig 9:**
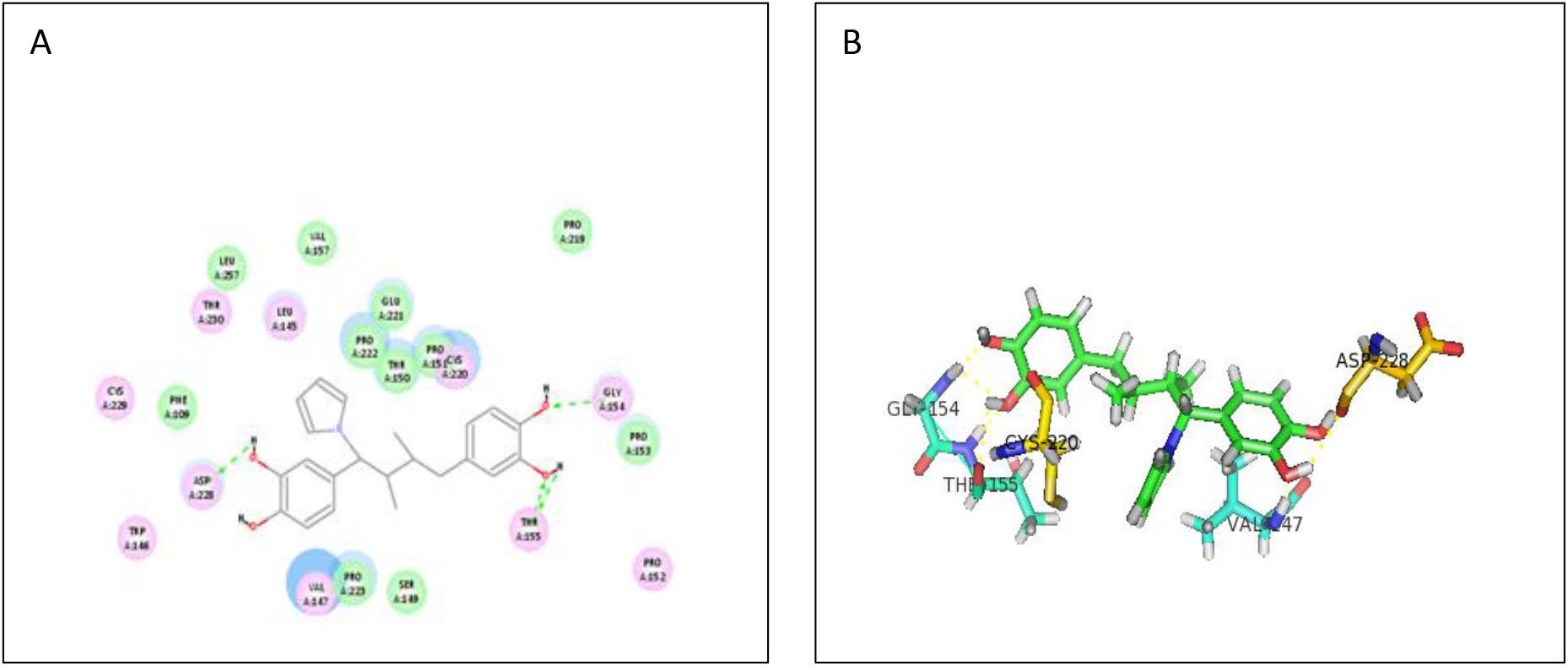
2D & 3D interaction map of Analog1 with target (501i)

**fig 10:**
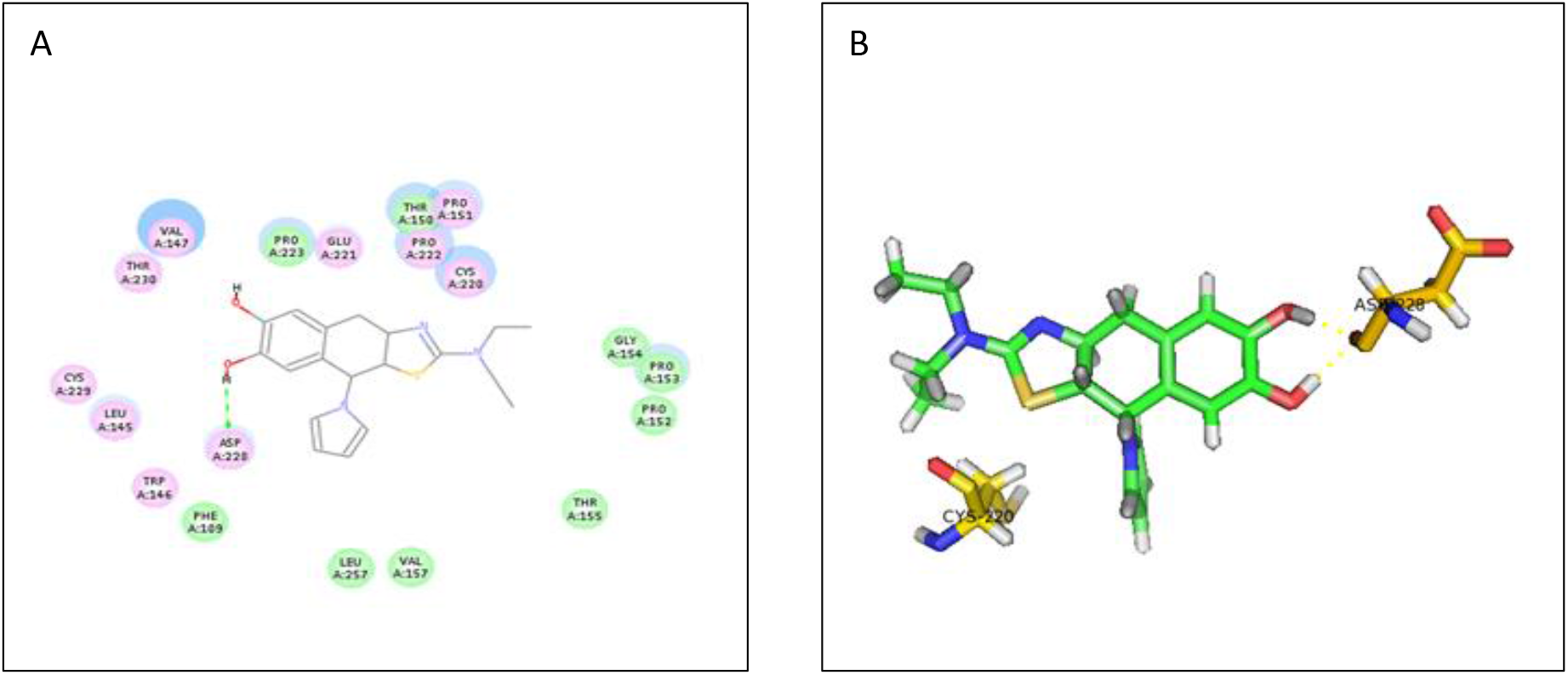
2D & 3D interaction map of Analog 2 with target (501i)

### 4.3 Interaction Comparison between Top compounds from the Library with Analog 1 & 2

Fragment-Based optimization produced 2 Analogs which were superimposed with Top binders MB710 and Phikan5196 (figure 11) and checked for key interactions, Position of pyrrole moiety as its acts as a Cys220 switch and N-ethyl group interactions and it was seen that the Analogs were able to maintain the interactions and cover the specific space in the cavity also the position of pyrrole group was accurate.

**fig 11:**
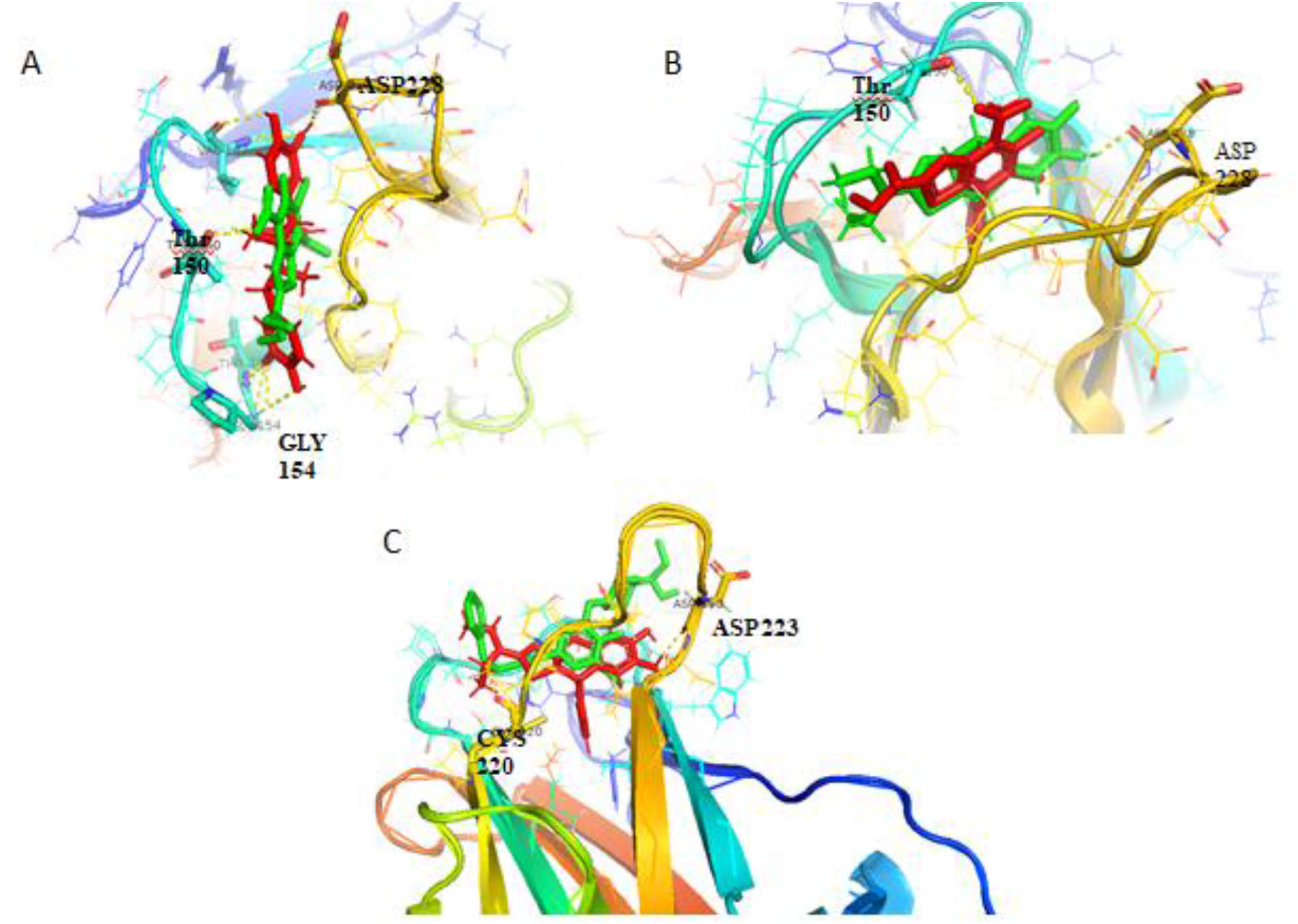
Superimposition of Docked pose of (A) MB710 and Analog 1 (B) MB710 and Analog 2 (c) Phikan5196 and Analog 2

**Fig 12:**
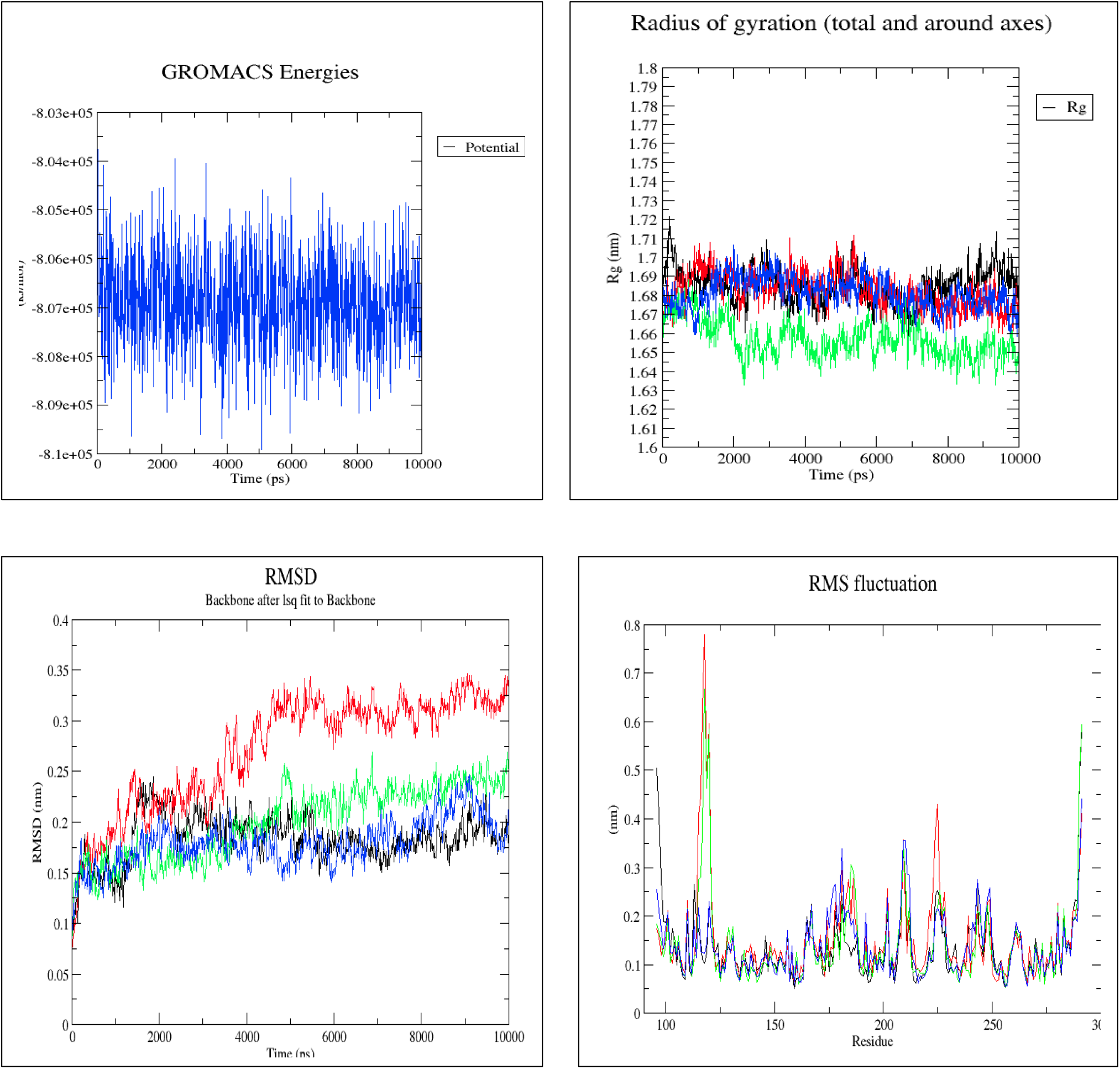
Simulation comparisons between MB710 (**Black**), Masoprocol (Red), Analog 1(green) and Analog 2(blue)

### 4.4 Molecular Dynamic simulations of Masoprocol Analogs in comparison with Masoprocol (parent compound) and MB710 (Reference Compound)

Substitution of Masoprocol raised 2 compounds Analog1 and Analog 2 and they got a Dock Score of 157.65 & 146.813 respectively which was higher than dock score of both reference and parent compounds also their simulation results in comparison with reference compound and parent compound showed that Analog 1 and Analog 2 were much more stable than the parent compound, RMSD curve of Analog1 is very close to that of reference compound and RMSD curve of Analog2 was overlapping and in some parts more stable than that of reference compound also RMSF curve from residue 222 to 230 (mutation domain) showed that Analog 2 stabilizes the mutation better than the reference compound (fig 11)

## 5. ADME analyses of the Analogs

Masoprocol analogs were submitted to the SwissADME server to compute physicochemical descriptors as well as to predict ADME parameters, pharmacokinetic properties, drug-like nature and medicinal chemistry friendliness of the designed analogs. The output from the SwissADME server predicts that both Analog1 and Analog2 have high Gastrointestinal absorption (pharmacokinetics), High Druglikliness and follow all the rules like Lipinski RO5, Veber rule, etc. and has Synthetic accessibility of 3.34 and 4.47 for Analog1 and Analog2 respectively (Table 6).

**TABLE 5:**
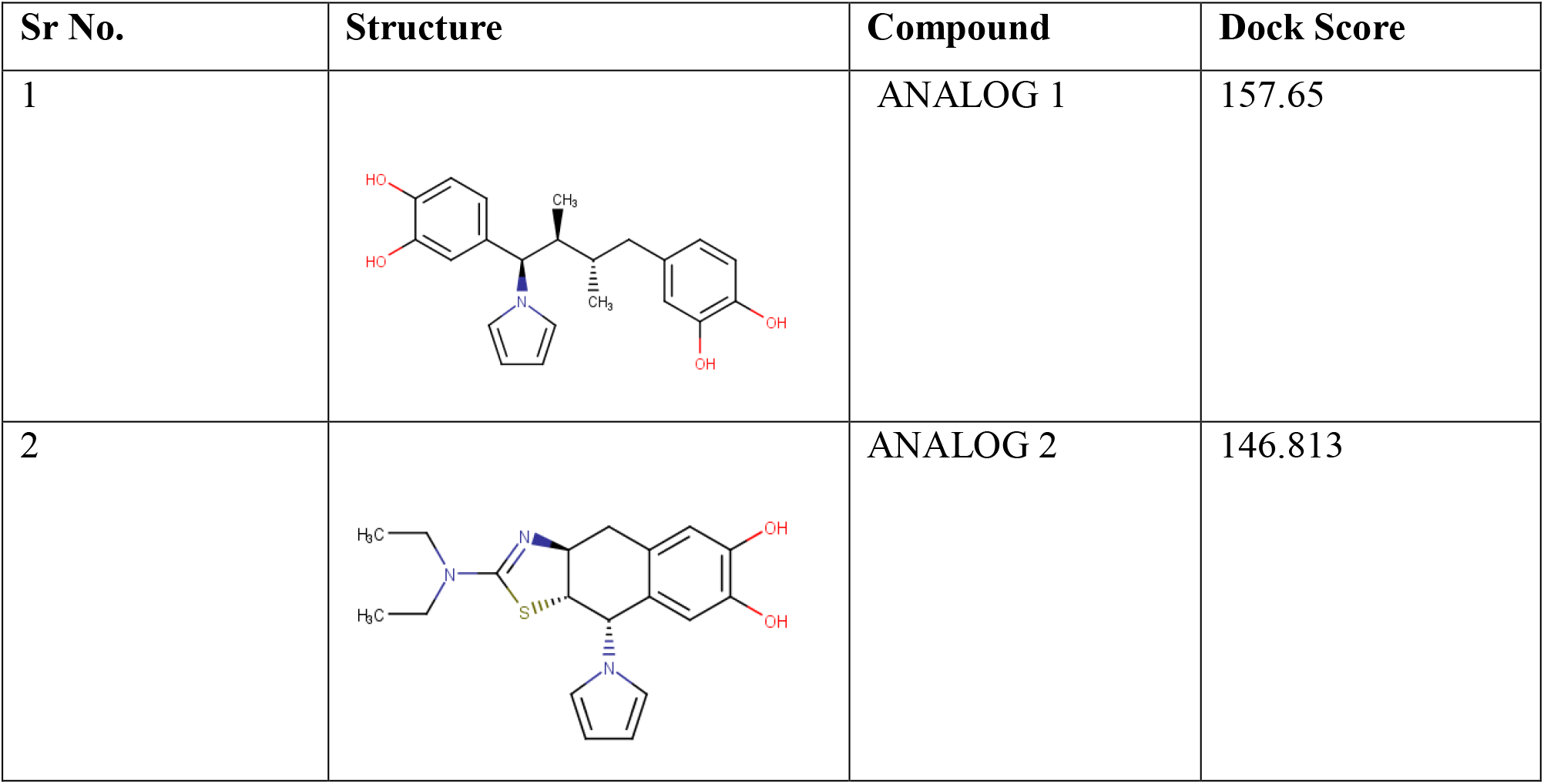
Docking and interaction studies of Masoprocol Analogues

| Sr No. | Structure | Compound | Dock Score |
| --- | --- | --- | --- |
| 1 |  | ANALOG 1 | 157.65 |
| 2 |  | ANALOG 2 | 146.813 |

**TABLE 6:** ADME and Drug like properties of Analog 1 and Analog 2

| Molecule | Physiochemical properties | Pharmacokinetics | Druglikeliness | Medicinal Chemistry |
| --- | --- | --- | --- | --- |
| Analog 1 | Formula<br>C<sub>22</sub>H<sub>25</sub>N<sub>04</sub> | GI absorption<br>High | Lipinski<br>Yes; 0 violation | LeadLikeness<br>No: Mw>350 |
| Analog 1 | Molecular weight<br>367.44g/mol | BBB permanent<br>No | Ghose<br>Yes | Synthetic<br>accessibility<br>3.34 |
| Analog 1 | Num heavy atoms<br>27 | P-GP substrate<br>Yes | Veber<br>Yes |  |
| Analog 1 | Num arom heavy atoms 17 | CYP1A2 inhibitor<br>No | Egon<br>Yes |  |
| Analog 1 | Fraction Csp3<br>0.27 | CYP2C19 inhibitor<br>Yes | Muegge<br>Yes |  |
| Analog 1 | Num rotatable bonds<br>6 | CYP3C9 inhibitor<br>No | Bioavailability<br>0.55 |  |
| Analog 1 | Num H-bond acceptors 4 | CYP2D6 inhibitor<br>Yes |  |  |
| Analog 1 | Num H-bond DONORS 4 | CYP3A4 inhibitor<br>Yes |  |  |
| Analog 1 | Molar refractivity<br>106.79 | Log Kp skin Permeation<br>-5.40 |  |  |
| Analog 1 | TPSA<br>85.85 A |  |  |  |
| **Molecules** | **Physiochemical properties** | **Pharmacokinetics** | **Druglikliness** | **Medicinal Chemistry** |
| Analog 2 | Formula<br>C19H23N3O2S | GI absorption<br>High | Lipinski<br>Yes; 0 violation | LeadLikeness<br>No: Mw>350 |
| Analog 2 | Molecular weight<br>357.47g/mol | BBB permanent<br>No | Ghose<br>Yes | Synthetic accessibility<br>4.47 |
| Analog 2 | Num heavy atoms<br>25 | P-GP substrate<br>Yes | Veber<br>Yes |  |
| Analog 2 | Num arom heavy atoms 11 | CYP1A2 inhibitor<br>No | Egon<br>Yes |  |
| Analog 2 | Fraction Csp3<br>0.42 | CYP2C19 inhibitor<br>No | Muegge<br>Yes |  |
| Analog 2 | Num rotatable bonds<br>4 | CYP3C9 inhibitor<br>No | Bioavailability<br>0.55 |  |
| Analog 2 | Num H-bond<br>acceptors 3 | CYP2D6 inhibitor<br>Yes |  |  |
| Analog 2 | Num H-bond<br>DONORS 2 | CYP3A4 inhibitor<br>No |  |  |
| Analog 2 | Molar refractivity<br>106.11 | Log Kp skin<br>permeation -<br>6.66 |  |  |
| Analog 2 | TPSA<br>85.29A |  |  |  |

Synthetic accessibility represents how easy or difficult is it to chemically synthesize the designed drug, SwissADME gives a score to the submitted drug which is in between 1 to 10 where 1 represents very easy to synthesize whereas 10 represents very difficult to synthesize

## 6. Conclusions

p53 is a homotetramer that specifically binds to its DNA consensus and attains structurally its most stable form. Y220C mutant p53 is unstable and degrades at physiological temperature and small molecules can be used to reactivate its activity. Masoprocol can be an important Lead for drug designing against the Y220C mutant p53 and more studies are required to design Masoprocol analog with higher potency.Analog1 and Analog2 show positive results in silico and specifically, Analog2 retains all the interactions until the end of simulations and shows a better result than the existing Reference compound MB710. Analog2 has the scaffold characteristics of MB710 along with the specific interaction with ASP-228 as present in Phikan5196 (Fig 10) which the second strongest binder and forms water-mediated H-bond with ASP-228 and this can be the reason behind its potency but further studies are required and Analogs are needed to be chemically synthesized and tested in cancer cell lines to validate the results in vitro.

## Abbreviations

FBHD: Fragment-based hybrid design
MD: Molecular Dynamic Simulations
RMSD: Root mean square deviation
RMSF: Root mean square fluctuation
SBDD: Structure-Based drug design
VS: Virtual Screening
QSAR: Quantitative structure-activity relationship
DBD: DNA binding Domain
OD: Oligomerization domain
CTD: C-terminal regulatory domain
ADME: Absorption Distribution Metabolism Excretion
SDS: Sodium Dodecyl Sulfate
PAGE: Polyacrylamide Gel Electrophoresis

## Acknowledgment

We thank Banaras Hindu University for its facilities. M R C thanks Department of Biotechnology, Government of India for scholarship and Centre for Bioinformatics Pondicherry University for providing Computational assistance.

